# Differential Immunoregulation by Human Surfactant Protein A Variants Determines Severity of SARS-CoV-2-induced Lung Disease

**DOI:** 10.1101/2024.09.11.612497

**Authors:** Ikechukwu B. Jacob, Akinkunmi O. Lawal, Salma S. Mahmoud, Emerson M. Kopsack, Erin S. Reynolds, Qinghe Meng, Hongkuan Fan, Paul T. Massa, Saravanan Thangamani, Hongpeng Jia, Guirong Wang

## Abstract

COVID-19 remains a significant threat to public health globally. Infection in some susceptible individuals causes life-threatening acute lung injury (ALI/ARDS) and/or death. Human surfactant protein A (SP-A) is a C-type lectin expressed in the lung and other mucosal tissues, and it plays a critical role in host defense against various pathogens. The human SP-A genes (*SFTPA1* and *SFTPA2*) are highly polymorphic and comprise several common genetic variants, i.e., SP-A1 (variants 6A^2^, 6A^4^) and SP-A2 (variants 1A^0^, 1A^3^). Here, we elucidated the differential antiviral and immunoregulatory roles of SP-A variants in response to SARS-CoV-2 infection *in vivo*. Six genetically-modified mouse lines, expressing both hACE2 (SARS-CoV-2 receptor) and individual SP-A variants: (hACE2/6A^2^ (6A^2^), hACE2/6A^4^ (6A^4^), hACE2/1A^0^ (1A^0^), and hACE2/1A^3^(1A^3^), one SP-A knockout (hACE2/SP-A KO (KO) and one hACE2/mouse SP-A (K18) mice, were challenged intranasally with 10^3^ PFU SARS-CoV-2 or saline (Sham). Infected KO and 1A^0^ mice had more weight loss and mortality compared to other mouse lines. Relative to other infected mouse lines, a more severe ALI was observed in KO, 1A^0^, and 6A^2^ mice. Reduced viral titers were generally observed in the lungs of infected SP-A mice relative to KO mice. Transcriptomic analysis revealed an upregulation in genes that play central roles in immune responses such as *MyD88*, *Stat3*, *IL-18*, and *Jak2* in the lungs of KO and 1A^0^ mice. However, *Mapk1* was significantly downregulated in 6A^2^ versus 1A^0^ mice. Analysis of biological pathways identified those involved in lung host defense and innate immunity, including pathogen-induced cytokine, NOD1/2, and Trem1 signaling pathways. Consistent with the transcriptomic data, levels of cytokines and chemokines such as G-CSF, IL-6 and IL-1β were comparatively higher in the lungs and sera of KO and 1A^0^ mice with the highest mortality rate. These findings demonstrate that human SP-A variants differentially modulate SARS-CoV-2-induced lung injury and disease severity by differentially inhibiting viral infectivity and regulating immune-related gene expressions.

## Introduction

COVID-19 is an infectious disease caused by severe acute respiratory syndrome coronavirus 2 (SARS-CoV-2) ^1,2^. Severe disease is characterized by a persistence of viral RNA and a robust influx of proinflammatory cytokines into the lungs, leading to acute lung injury (ALI) and acute respiratory distress syndrome (ARDS) ^1,2^. Barely four years since first reported, there have been more than 7 million deaths due to COVID-19 globally ^3^. Although most people recover from acute infection without requiring hospitalization, some develop severe morbidity and mortality even in the absence of known risk factors ^4^. While Genome-Wide Association Studies (GWAS) have widely demonstrated differential disease outcomes in the general population, with multiple studies implicating genetic variations in host innate immunity, the reasons for the differential disease phenotypes are still unclear ^4–8^.

Human surfactant protein A (SP-A) is a C-type lectin (collectin) expressed by alveolar epithelial type 2 cells (AT2) of the lungs and other mucosal tissues. It plays critical roles in innate host defense, immunomodulation, and surfactant physiology in the lung and other mucosal surfaces ^9^. SP-A can bind glycoconjugates on microbes and interact with host cell receptors to inhibit infectivity and maintain lung homeostasis ^9^. Our data showed that SP-A can bind to SARS-CoV-2 spike protein (S), inhibit viral infectivity in lung epithelial cells, and protect against severe lung injury in humanized transgenic (hTG) mice^10^. Other investigators have previously demonstrated the importance of collectins in attenuating viral infectivity and disease severity in the context of SARS-CoV-2, Influenza A virus (IAV), Human Immunodeficiency Virus (HIV), and Respiratory Syncytial Virus (RSV) infections ^10–19^.

The human SP-A protein is encoded by two highly functional and polymorphic genes (*SFTPA1*: SP-A1 and *SFTPA2*: SP-A2). Several genetic variants (alleles) of these genes have been identified in the general population, such as *SP-A1* (variants: 6A^2^, 6A^4^) and *SP-A2* (variants: 1A^0^, 1A^3^) ^20^. Although there are only a few amino acid differences between SP-A1 and SP-A2 variants, SP-A2 variants have a higher capacity to bind carbohydrates, enhance phagocytosis of invading pathogens by macrophages and induce higher proinflammatory cytokine production in immune cells under various stressed conditions, indicative of the potential of SP-A variants to differentially regulate immune function in the context of infection ^21–24^. Furthermore, SNPs resulting in an amino acid change from arginine to tryptophan in the 6A^4^ variant have been demonstrated to alter its structure and function, enhancing its susceptibility to pulmonary diseases, including tuberculosis ^20,25–27^. Previous population-based studies found that differences in collectin genes such as those of the Mannose-binding lectin (MBL), surfactant protein D (SP-D), and SP-A play important roles in disease severity following viral and bacterial infections ^28–32^; while dysfunctions in surfactant-related genes were shown to associate with severe COVID-19 ^33^. Importantly, recent GWAS studies have strongly implicated SP-A and SP-D genes in the differential COVID-19 susceptibility and severity observed in the human population ^34,35^. Although the latter studies point to the potential host defense role of SP-A genes during infection, mechanistic pre-clinical studies of variant-specific functions of human SP-A genes in response to SARS-CoV-2 infection are lacking. Therefore, an adequate animal model to study the roles of SP-A genetic variants on host defense and lung function following the SARS-CoV-2 challenge is vital.

In this study, we have generated and characterized double-humanized transgenic (double-hTG) mice expressing both hACE2 (SARS-CoV-2 cognate receptor) and single genetic variants of human SP-A (6A^2^, 6A^4^, 1A^0^ or 1A^3^). Using these mice, we explored whether there are differences in COVID-19 severity (mortality and ALI) and the underlying mechanisms of disease modulation by SP-A genetic variants post-SARS-CoV-2 (Delta variant) challenge. Our data revealed distinct antiviral and immunomodulatory roles of SP-A genetic variants in COVID-19 symptomology among hTG mouse lines. We observed two different patterns of disease severity based on animal mortality and ALI severity. First, we observed that 6A^2^ and 1A^3^ mice are relatively more protected against SARS-CoV-2-induced mortality compared to SP-A KO and 1A^0^ mice; second, 6A^4^ and 1A^3^ mice developed moderate ALI following SARS-CoV-2 infection. Essentially, we observed that in response to SARS-CoV-2 infection, SP-A genetic variants differentially modulate severe lung injury and survival rate by differentially regulating immune genes and inflammatory cytokine production locally and systemically. The data support ongoing efforts at producing personalized surfactant-based therapies for COVID-19 patients ^36,37^.

## Materials and Methods

### Mouse Models

Mice breeding and maintenance were carried out at SUNY Upstate Medical University’s animal core facility. The animals were housed in a temperature-controlled room at 22 °C under specific pathogen-free conditions. Mice between 8-16 weeks of age of both sexes were used in this study. The animal experiments were approved by SUNY Upstate Medical University Institutional Animal Care and Use Committee and conducted according to the National Institutes of Health and ARRIVE guidelines on using laboratory animals. The initial K18 (hACE2/mSP-A) mice were obtained from Jackson Laboratories (Bar Harbor, ME); these mice are susceptible to SARS-CoV-2 infection since they express the hACE2 transgene. To examine the role of human SP-A variants in the context of SARS-CoV-2 infection, we generated transgenic mice that carry both hACE2 and SP-A transgenes (double humanized transgenic or double-hTG) in which the hACE2 and individual single-gene variants of SP-A1 (6A^2^ and 6A^4^) and SP-A2 (1A^0^ and 1A^3^) are expressed and the mouse SP-A gene has been deleted by breeding with previously characterized hTG SP-A mice ^38^. To do this, we initially crossed K18 mice that are hemizygous for the hACE2 transgene with our SP-A knockout (KO) mice to remove the mSP-A gene. Genotyping was performed after each filial generation. Subsequently, we crossed the hACE2/SP-A KO mice with mice carrying individual variants of either SP-A1 (6A^2^ or 6A^4^) or SP-A2 (1A^0^ or 1A^3^) gene to generate a double-hTG mouse model with hACE2 and single-gene variants of human SP-A. We examined the presence of hACE2 and human SP-A genes in the mice by PCR genotyping and analyzed SP-A expression in the lung by immunoblotting. The double-hTG mice (hACE2/6A^2^ (6A^2^), hACE2/6A^4^ (6A^4^), hACE2/1A^0^ (1A^0^) and hACE2/1A^3^ (1A^3^) alongside hACE2/SP-A KO (KO) and hACE2/mSP-A (K18) mice were subsequently used for SARS-CoV-2 (Delta) challenge studies.

### Mouse Infection and Sample Collection

SARS-CoV-2-and mock-challenged mice (6 groups: 6A^2^, 6A^4^, 1A^0^, 1A^3^, K18, and KO) were anesthetized with isoflurane and infected intranasally (i.n.) with 30 µl (15 µl/nose) of virus solution containing 1× 10^3^ PFU of SARS-CoV-2 in 1X MEM media. Control (Sham) mice were inoculated with 30 µl of 1X MEM. After viral infection, mice were observed daily for morbidity (body weight) and mortality (survival). Mice showing >25% decrease in their initial body weight were defined as reaching the experimental endpoint and euthanized. Mice were sacrificed by anesthesia and exsanguination on days 2, 4, and 6 post-infection (pi) to collect lung samples for viral load analysis by RT-qPCR, immunohistochemistry (IHC), and plaque assay. Cytokine and gene-expression changes in the lung were analyzed, and systemic cytokines in sera were also determined as described previously ^39,40^.

### Tissue Processing

Mice were anesthetized with isoflurane and exsanguinated. Then, we perfused each mouse with 3 mL NBF. Then the lungs were inflation-fixed by tracheal instillation of 1 mL NBF. Mouse whole bodies were dissected along the abdomen (n = 7-11 mice /group) and fixed in NBF before histological and immunohistochemical analyses. Lungs from a subset of mice were harvested (n = 6-7 mice/group) and frozen immediately at -80 ^°^C for further processing. The lungs were weighed and homogenized in cold PBS and then centrifuged for 5 mins at 21130x g and supernatants were collected for viral load and serological analyses using RT-qPCR, plaque assay, and ELISA as described previously ^10,40^.

### Histopathological Analysis

Fixed lung tissues were embedded as previously described ^40^. 5 µm sections of individual tissue samples were cut and stained with Hematoxylin and Eosin (H&E) and then microscopically examined. Lung injury was scored using a 0-2 scale by counting the number of neutrophils in the alveolar space, neutrophils in the interstitial space, presence of hyaline membranes, proteinaceous debris filling the airspaces, and alveolar septal thickening as described by ^41^. Histopathological analyses were blinded and carried out by two independent pathologists.

### Immunohistochemistry

Samples were analyzed by IHC as described previously by ^40^, with some modifications. Sections were deparaffinized in an oven for 1 h at 60°C and then rehydrated sequentially using 100%, 95%, 80%, 70% and 50% ethanol. The sections were rinsed twice in double distilled water and boiled in 10 mM citrate (pH 6.0) for 9 mins for antigen retrieval followed by cooling at room temperature for 20 mins. The sections were blocked with 10% BSA in PBS for 1 h at room temperature to prevent nonspecific background staining. This was followed by overnight incubation of the slides at 4°C with SARS-CoV-2 nucleocapsid protein (NP) rabbit antibody (cell Signaling, #33336S; 1:100). The next day, after several washes, the sections were incubated with biotinylated goat anti-rabbit antibody for 1 h (1:200), followed by staining using ABC/HRP complex (Vectastain ABC kit, peroxidase rat IgG PK-4004, and peroxidase rabbit IgG PK-4001; Vector Laboratories). Stained sections were visualized using 3’3-diaminobenzidine (DAB) (SK-4100; Vector Laboratories) and counterstained with hematoxylin (H-3404-100) (QS counterstain). Images were acquired using the Nikon Eclipse TE2000-U microscope (Nikon Corporation, Tokyo, Japan) and the number of SARS-CoV-2 NP-stained lesions was counted using Image J software.

### Western Blotting Analysis

Lung tissues from each group were homogenized in RIPA buffer (ThermoFisher Scientific, Rockford, IL) containing a cocktail of protease and phosphatase inhibitors (Roche, Indianapolis, IN). Total protein concentration in lung homogenates was evaluated by the microBCA assay kit (ThermoFisher Scientific), following the manufacturer’s instructions. Thirty micrograms of total protein were subjected to SDS-PAGE on a 10% gel in reducing conditions and transferred to PVDF membranes (Bio-Rad). To block nonspecific interactions, the blots were incubated in a buffer with 5% non-fat milk for 30 mins before further incubation with SP-A polyclonal antibody (1:1000). As a loading control, blots were re-probed with β-actin (1:1000, ab-16039, Abcam, MA, USA). Membranes were developed and observed by incubating with goat anti-rabbit IgG HRP-conjugated secondary antibody (Bio-Rad) and by using the ECL Western Blotting substrate as described previously ^38^.

### Measurement of viral loads

Total RNA was isolated from homogenized lung tissues using the Quick-RNA extraction miniprep kit (# R1055 Zymo Research, CA, USA) following the manufacturer’s instructions and the concentration of RNA was measured using the nanodrop machine (Thermo Scientific). Real-time qPCR (RT-qPCR) was performed using the AB StepOnePlus Detection System (Applied Biosystems, Foster City, CA) using the one-step kit RT-PCR Master Mix Reagents (#64471423, Biorad), prepared according to the manufacturer’s instructions. Purified RNA (100 ng) was added into a 10 µl total volume of real-time PCR mix buffer containing forward/reverse primer pairs (forward, AGCCTCTTCTCGTTCCTCATCAC; reverse, CCGCCATTGCCAGCCATTC; each 500 nM) targeting SARS-CoV-2 N1 gene and a probe (250 nM, FAM) and other reagents provided by the manufacturer. The one-step RT-qPCR was performed through a cycle of reverse transcription at 55°C for 10 mins followed by 40 cycles of amplification at 95°C for 3 mins, 95°C for 15 s, and 55°C for 1 min. SARS-CoV-2 copy numbers in lung tissues were quantified using SARS-CoV-2 RNA standards as described by ^42^.

Viral titer was also quantified using supernatants from lung homogenates by plaque assay. In brief, Confluent monolayers of Vero E6 in 24-well plates (1 × 10^5^ cells/well seeded) were infected with 10-fold serial dilutions of supernatants from each mouse group. The cells were cultured for 1 h with intermittent rocking for viral adsorption. After which, the unbound virus was removed by washing with PBS, overlayed with 2% methylcellulose and cultured for another three days at 37°C in a humidified incubator. After plaque development, cells were fixed with 10% formalin for 60 mins and stained with 0.05% (w/v) crystal violet in 20% methanol and plaques were quantified as PFU/ml.

### Transcriptomics Analyses

The expressions of 84 innate and adaptive immune genes in the lungs of SARS-CoV-2 infected mice and Sham were determined using RT^2^ Profiler^TM^ PCR array kit (Cat#: PAMM-052ZC-24, Qiagen) ^39^. In brief, total RNA was isolated from homogenized lung tissues using the Quick-RNA extraction miniprep kit (# R1055 Zymo Research, CA, USA) following the manufacturer’s instructions and RNA concentration and purity was determined by spectrophotometry using the nanodrop machine (Thermo Scientific). cDNA was synthesized from 500 ng of total RNA using the iScript Reverse Transcription Supermix for RT-qPCR (Cat#: 1708841, Biorad). Real-time PCR was performed using the RT^2^ Profiler^TM^ PCR array system following the manufacturer’s recommendation in the AB StepOnePlus Detection System (Applied Biosystems, Foster City, CA). The levels of expression of the mRNA of each gene in the different groups were normalized to the housekeeping gene (*HSP90*). Data was exported into an Excel spreadsheet and analyzed with Qiagen’s PCR analysis web-based software (GeneGlobe Data Analysis Center). Relative gene expression changes were calculated by the 2^−(averageΔΔCt)^ method. Fold change (FC) values are compared relative to the sham group or compared between two infected groups. Gene expression FCs are considered significant when FC > 2.0, p-value > 0.05). We further used specific primers to determine the expression of some significantly upregulated genes in infected hTG mice by RT-qPCR and then normalized to Gapdh expression (Extended Table 1 shows the primer list).

### Signaling Pathway Association Analysis

Differentially expressed genes (DEGs) were functionally analyzed using the Qiagen’s Ingenuity Pathway Analysis (IPA, QIAGEN, Redwood City, CA) software and the analyses included canonical pathway analysis, gene networks and upstream regulators. Canonical pathways were scored by analyzing the ratio of the number of genes that map to a specific pathway in the IPA database. The observed gene networks show functional relationships among DEGs based on database associations. IPA was used to gain additional insights into the immunoregulatory roles of each SP-A humanized mouse in response to SARS-CoV-2 infection.

### ELISA of Lung Tissue Cytokines

Levels of TNF-α (Lot# 227846-003), IL-1β (#372944-002), IL-6 (#330384-008), IL-10 (#355429-004), MCP-1 (#175884-002) and IFN-γ (#300277-008) in mouse lungs were measured by enzyme-linked immunosorbent assay (ELISA) ^43^. All measurements were in duplicate with kits according to the manufacturer’s instructions and the fold changes represented in the heat map among the infected mouse groups are relative to those observed in the mock-infected group.

### Serum Cytokine and Chemokine Analysis: Multiplex Cytokine Assay

Inflammatory cytokines and chemokines in sera were analyzed using a Bioplex Mouse cytokine 23-plex panel magnetic bead Luminex assay (Biorad, 23-plex, Group 1, #64436152) following the manufacturer’s instructions. Twenty-five microliters of serum samples were diluted in 75 µl Bio-plex sample diluent and incubated with magnetic capture beads. Then, the plate was washed, and detection antibodies were added followed by streptavidin incubation. The analytes were measured using a Luminex 200 Analyzer (Luminex) and quantitated using a standard curve. Samples from each mouse were analyzed in duplicate (50 µl per well). All immunoassays were performed in the BSL-3 facility and fold changes represented in the heat map among the infected mouse groups are relative to those observed in the mock-infected group.

### Statistical analysis

All experimental data are presented as mean ± standard error and statistically analyzed using GraphPad Prism 8.0 (GraphPad Software, San Diego, CA, USA). Comparisons between two independent groups were performed using Student’s *t*-test or multiple groups using one-way ANOVA. Survival analysis was performed with Kaplan-Meyer survival curves and evaluated statistically by the log-rank test. Results were considered statistically significant when P<0.05.

## Results

### Generation and Characterization of Double-hTG Mice

To study the differential role of SP-A genetic variants in response to SARS-CoV-2 infection in the lungs, we generated double-hTG mice expressing both hACE2 (SARS-CoV-2 cognate receptor) and individual SP-A genetic variants derived from either SP-A1 (6A^2^ and 6A^4^) or SP-A2 (1A^0^ and 1A^3^) genes. This was done by crossing K18 mice expressing hACE2 and mSP-A with SP-A KO mice to generate mice that are positive for hACE2 and eliminate the mSP-A gene (Fig 1A). These KO mice, positive for hACE2 (hACE2/KO) were then bred with our previously characterized SP-A hTG mice carrying single-gene variants 6A^2^, 6A^4^, 1A^0^, or 1A^3^ to generate mice that carry both hACE2 and individual SP-A variant. The representative genotyping analysis of hACE2, SP-A1, SP-A2, and mSP-A genes from each mouse line is shown in Fig 1B. As expected, all the mice expressed hACE2, but KO mice had no human or mouse SP-A by PCR analysis. SP-A protein expression in the lungs of KO, K18, 6A^2^, 6A^4^, 1A^0^ and 1A^3^ mice was analyzed by Western blotting using an anti-SP-A antibody. Except for KO mice, SP-A was expressed in all SP-A hTG mice as expected (Fig 1C).

**Fig. 1.**
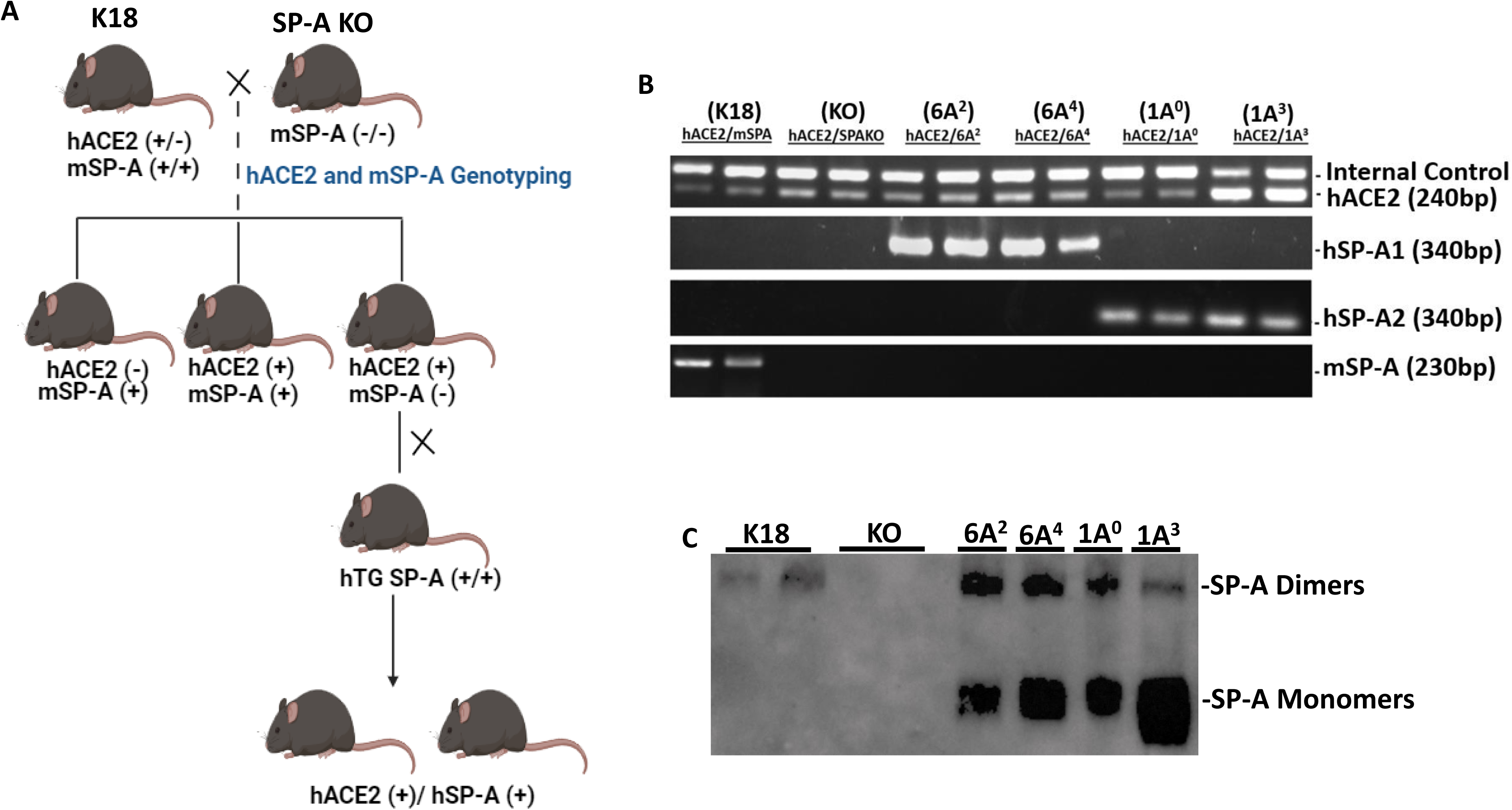
Characterization of double-humanized transgenic mice. Double-humanized transgenic (double-hTG) mice expressing both hACE2 and human SP-A1 (6A^2^ or 6A^4^), or SP-A2 (1A^0^ or 1A^3^) genetic variants were generated and characterized using the breeding strategy shown in Panel A. We also generated mice with hACE2 but SP-A deficient (hACE2/KO) and those with hACE2 and mSP-A (K18). (A): Breeding scheme of double-hTG mice generated by crossing K18 mice with SP-A KO mice to generate hACE2/KO mice. hACE2/KO mice were subsequently crossed with previously characterized hTG SP-A mice. (B): Genotyping analysis of double-hTG, KO and K18 mice by PCR. C: Western blotting analysis of SP-A expression in the lung of double-hTG mice showing hSP-A expressions in the lungs of double-hTG and K18 but not in mice deficient in SP-A (KO).

### Significant Body-Weight Loss and higher Mortality rate in SP-A KO and 1A^0^ Mice

Several population-based studies have shown that human SP-A variants may be either detrimental or protective depending on the disease ^20,44^. Thus, to gain insights into the roles of human SP-A variants with regard to SARS-CoV-2-induced morbidity and mortality, we infected four different SP-A hTG mice: 6A^2^, 6A^4^, 1A^0^ and 1A^3^, one K18 and one KO mice (Fig. 2A). We observed that weight loss was particularly severe in KO mice and those carrying the 1A^0^ variant, particularly on days 5 and 6 post-viral challenge. On day 6 pi, KO, 1A^0^ and 6A^4^ mice had lost more than 13%, 12%, and 10% of their initial weights, respectively (Fig 2B). Although body-weight changes were observed early on day 3 pi. in KO and not in 1A^0^ mice, there was a drastic decrease in body weight in 1A^0^ mice by days 5 and 6 pi., suggestive of a differential response of the different SP-A variants to viral-challenge. Importantly, compared to KO mice on day 6 post challenge, mice carrying the 6A^2^ and 1A^3^ variants did not significantly lose body weight (P<0.05). The results until day 14 pi. showed that most of the mouse groups except KO, gradually regained their body weights until the end of the study. However, none of the KO mice survived beyond day 12 pi. Other clinically relevant characteristics observed in infected mice were ruffled fur, lethargy/unresponsiveness to stimuli, shivering, hunched posture, mucus around eyes (observed in the most severe mice), and labored breathing.

**Fig. 2.**
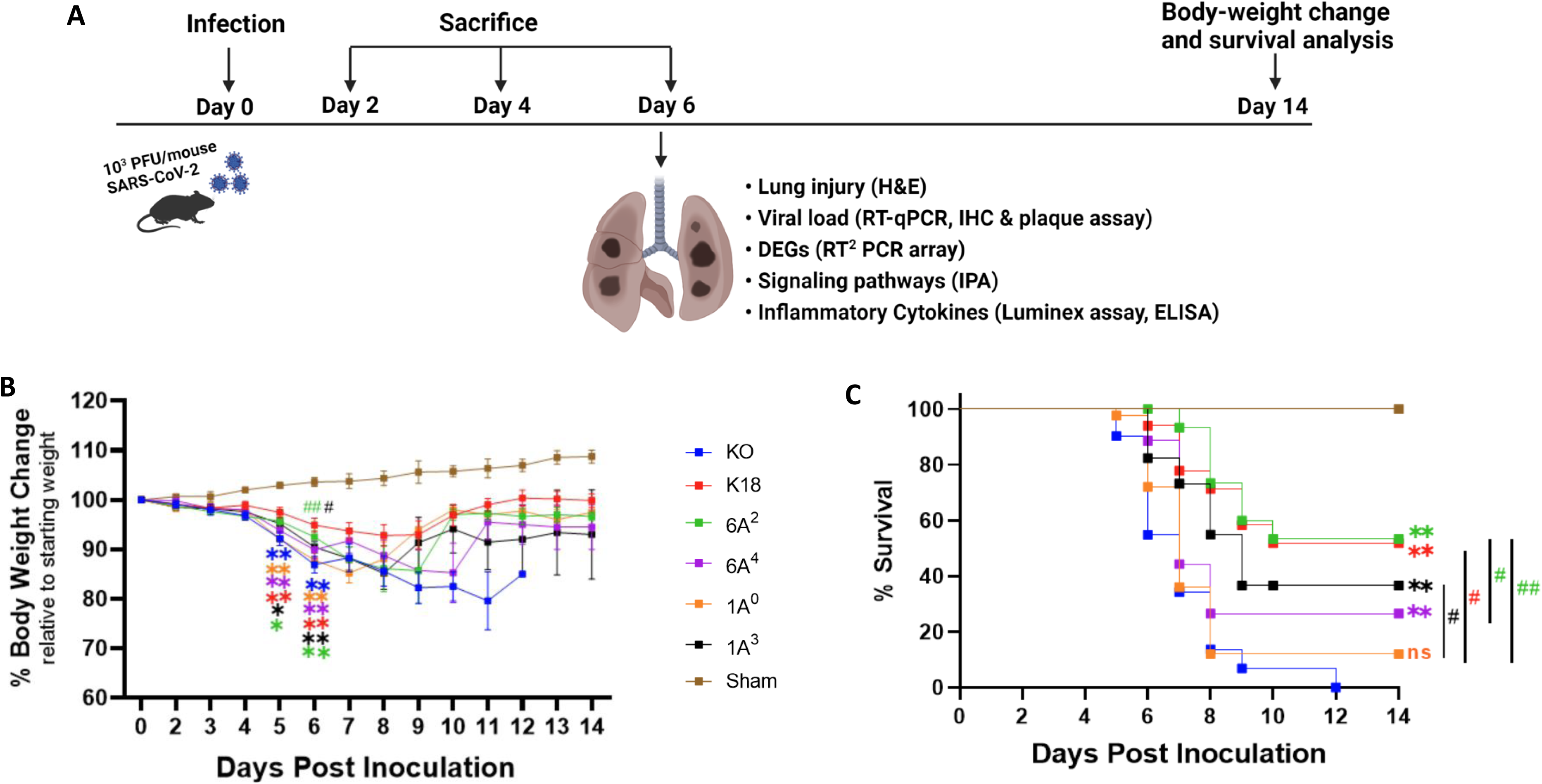
Body-weight loss and mortality in infected humanized mice. A total of six transgenic mouse lines, four of them carrying hACE2 alongside distinct human SP-A variants, i.e. hACE2/6A^2^ (6A^2^), hACE2/6A^4^ (6A^4^), hACE2/1A^0^ (1A^0^), and hACE2/1A^3^ (1A^3^), and the other two lines, i.e. hACE2/SP-A KO (KO) and hACE2/mSP-A (K18), were infected intranasally with 10^3^ PFU/mouse of SARS-CoV-2 (Delta variant) or Sham (30 µl of MEM medium) on day 0; and animals were monitored daily for survival and body weight changes until 14 dpi or sacrificed at the indicated timepoints as shown in Panel A. Panels B and C showed the body-weight changes and animal survival of each group, respectively. (n= 18-20 mice/group). Symbols in (B) represents mean ± SEM. \**P*< 0.05, \*\**P*< 0.01 (relative to sham), #<0.05, ##<0.01 (relative to infected KO), n=5 independent experiments. NS = not significant. Color corresponds to each mouse line.

On day 5 pi, some of the 1A^0^ and KO mice had reached mortality as defined by a loss of >25% of starting weight and were euthanized as a humane experimental endpoint (Fig. 2C). By 6 dpi, about 45% of KO and 27% of 1A^0^ mice had died or reached the study endpoint for euthanasia. Strikingly, 100% survival was observed in mice carrying the 6A^2^ variant, closely followed by 88% and 82% survival observed in 6A^4^ and 1A^3^ mice, respectively, on day 6 pi. By day 14 pi. the results showed that mice carrying the 6A^2^ and 1A^3^ variants also had the highest survival rate (53.3% and 36.6% respectively) compared to 6A^4^ and 1A^0^ mice (26.6% and 12% respectively). KO mice, however, did not survive beyond day 12 pi. These findings suggest that mice carrying SP-A2 (1A^0^) genetic variant and those deficient in SP-A had more severe morbidity and higher mortality rates compared to those carrying SP-A1 (6A^2^ and 6A^4^) and SP-A2 (1A^3^).

### Human SP-A Variants Differentially Attenuate SARS-CoV-2-Induced ALI in Mice

As an important host defense protein and innate immune modulator, variations in the human SP-A gene have been associated with differences in susceptibilities and severities to acute and persistent pulmonary diseases ^45–47^. Therefore, we examined ALI features in mice carrying individual SP-A variants following SARS-CoV-2 infection. Lungs were collected on day 6 pi.; and the severity of lung injury in our mouse groups was evaluated in H&E-stained sections as recommended by the American Thoracic Society ^41^. In this study, we observed the presence of inflammatory cells in the alveolar space and interstitial (black arrows, Fig 3A and Extended Fig 1), proteinaceous debris within the airways, and a thickening of alveolar walls resulting from a loss of alveolar integrity and fluid accumulation (red arrows, Fig 3A) in infected lungs. As expected, none of these characteristics was found in the lung of sham mice (Fig. 3A).

**Fig. 3.**
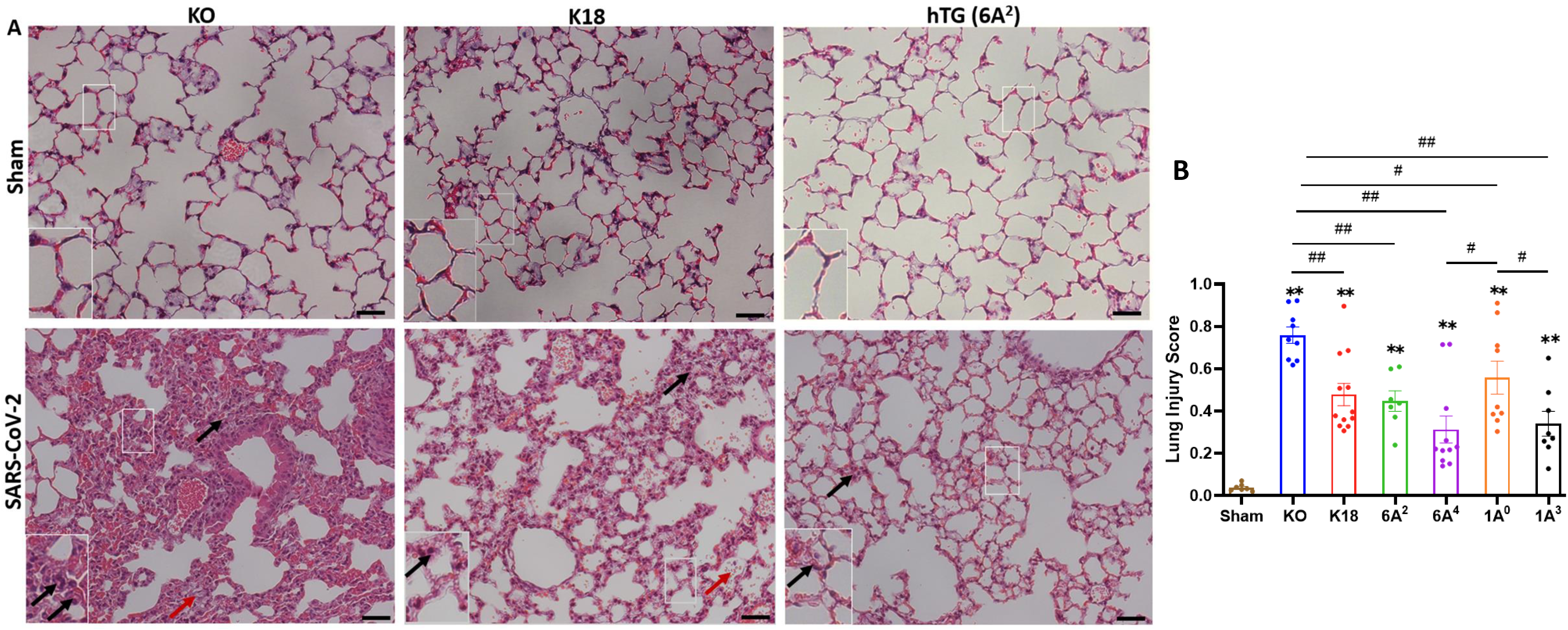
Differential lung injury after SARS-CoV-2 infection of humanized mouse lines. (A): Representative histological images of lungs from each mouse group on day 6 p.i. SARS-CoV-2 induced a more severe lung damage in KO mice characterized by an infiltration of inflammatory cells in alveolar spaces and interstitial (black arrows), accumulation of proteinaceous debris, and edema leakage of protein-rich fluid in the alveolar space (red arrows) compared to K18 and hTG SP-A1 6A^2^ (representative images are shown in Panel A). (B): Semi-quantitative histological assessment of lung injury from each mouse group showed significantly higher scores of lung injury in all infected groups compared to sham group, and differential lung injury scores were observed in mice carrying unique SP-A genetic variant. Symbols in (B) represents mean ± SEM of individual animals. \**P*< 0.05, \*\**P*< 0.01 (relative to sham), #<0.05, ##<0.01 (between two infected mouse groups from 5 independent experiments). Scale bars= 50 µm.

Interestingly, the markers of severe ARDS were more overt and severe upon scoring in KO mice relative to other groups, suggestive of the role of SP-A in host defense and maintaining alveolar integrity post-SARS-CoV-2 infection (KO vs hTG, P<0.01). However, a comparison of lung injury scores among the variants showed that 6A^4^ and 1A^3^ mice had the most attenuated ALI compared to those carrying 6A^2^ and 1A^0^ variants (the order of ALI score: 1A^0^>6A^2^>1A^3^>6A^4^) (Fig 3B). Further analysis of SP-A1 genetic variants showed that lung injury score trended higher in 6A^2^ relative to 6A^4^ mice while a comparison of SP-A2 gene-specific variants revealed a significantly higher lung injury score in 1A^0^ compared to 1A^3^ mice. These observations suggest that human SP-A variants 6A^4^ and 1A^3^ may be more protective against SARS-CoV-2-induced ALI compared to 6A^2^ and 1A^0^. Together, the results highlight two distinct outcomes of SARS-CoV-2-induced pathology in our experimental mouse groups based on body-weight changes and survival, and lung injury scores.

### Human SP-A Variants Differentially Attenuate SARS-CoV-2 Load in Mice

Given that viral burden has been associated with COVID-19 severity, we next quantified and compared viral load 6 dpi in infected mouse lungs from each group by RT-qPCR, IHC, and plaque assay ^48,49^. IHC staining of viral N proteins revealed higher levels of viral antigens in the alveolar septum and mononuclear cells in the lungs of SP-A deficient mice relative to the other hTG mice on day 6 pi (P<0.01). However, upon quantification, we observed comparatively reduced N proteins in the lungs of mice carrying the 6A^2^ and 1A^0^ variants on Day 6 p.i. (Fig 4A and 4B). Relative to KO mice with the highest viral genome copy numbers, the other groups (except K18) had significantly reduced copy numbers (p<0.05). Among the variants, the 1A^3^ variant had the highest average viral RNA copies (Log_10_ 8.6) (the order of viral RNA copies: 1A^3^>6A^4^>6A^2^>1A^0^) (Fig 4C). Furthermore, on day 6 pi, we were only able to consistently determine infectious viral particles in KO, K18, and 1A^3^ mice (∼10^3^ PFU/mL) while most other SP-A hTG lines had cleared the virus and infectious virus was therefore not observed at the limit of detection (LOD) (Fig 4D). To address the innate antiviral role of human SP-A, infectious virus titer was quantified in lungs collected on day 2 pi since there was viral clearance in most SP-A hTG lungs by day 6 pi. We observed a significant decrease in viral titer in K18, 6A^2^, and 1A^0^ mice relative to KO mice, with 1A^0^ mice having the lowest viral burden in the lungs; suggestive of the more profound antiviral role of 1A^0^ and 6A^2^ variants in response to SARS-CoV-2 infection *in vivo* (Fig 4D). Further analysis of lungs collected on day 4 pi showed a decline in viral titer compared to those on day 2 pi. The results suggest that viral load contributes to disease severity in mice given the attenuated ALI and death recorded in mice carrying human SP-A genetic variants relative to KO. SP-A may attenuate SARS-CoV-2 severity through its inhibitory effect on virus infectivity. However, 1A^0^ mice with the most severe mortality and ALI had the lowest viral burden in lungs at the time points studied; suggestive of the involvement of other distinct functions of SP-A in controlling disease severity besides its antiviral function.

**Fig. 4.**
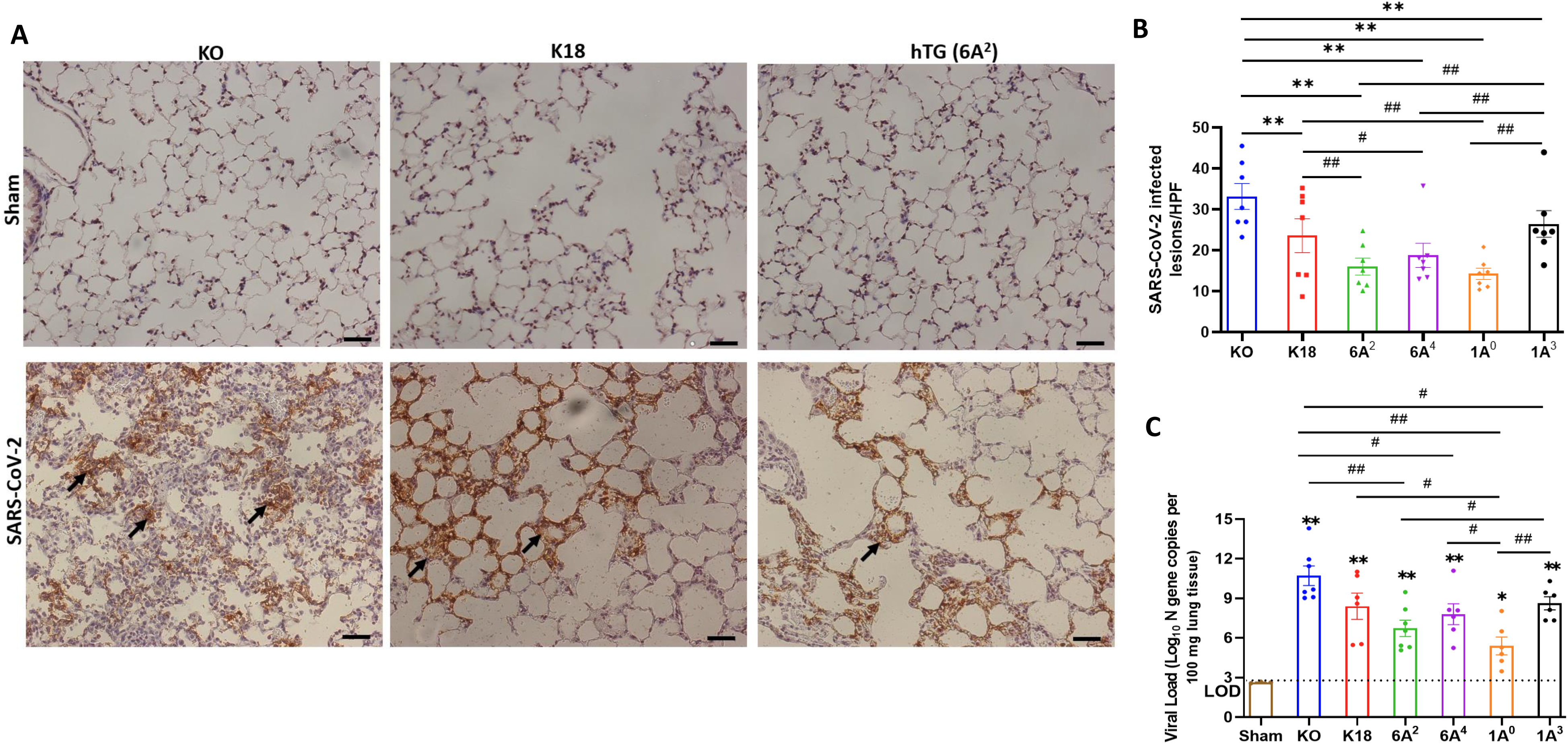
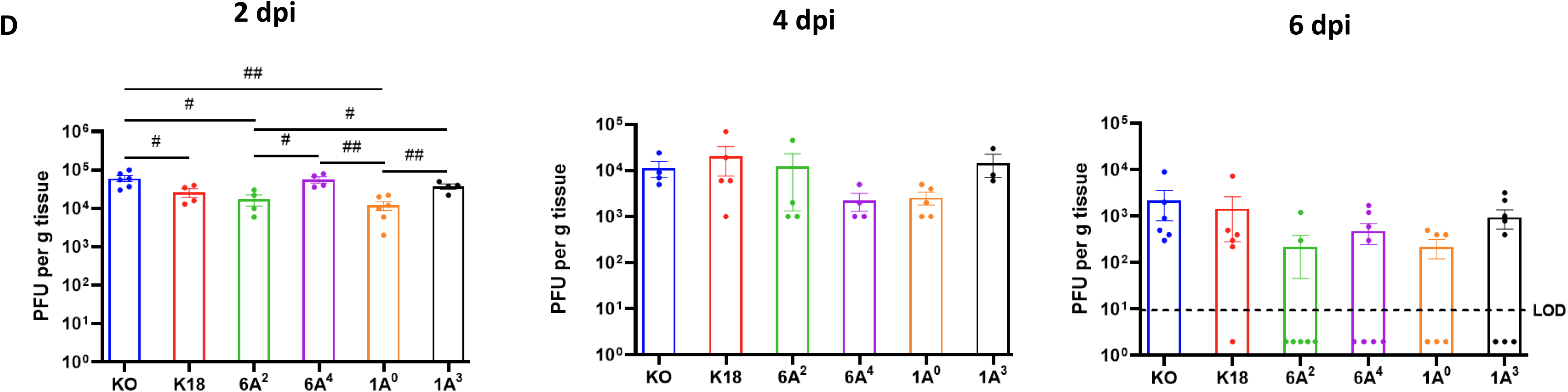
Differential viral load in the lung of SARS-CoV-2 infected humanized mouse lines. Viral load in the lungs of SARS-CoV-2-infected mice were determined using IHC, RT-qPCR and plaque assay. (A): IHC analysis was performed for SARS-CoV-2 nucleocapsid protein (NP) in the lungs of infected and sham groups on day 6 pi. Viral antigen was observed in all infected groups but not in sham. The arrows indicate the presence of viral antigens in the alveolar septum and cells. (B): Quantification of NP-positive lesions in each lung section reflected a significantly higher number of viral antigens in infected KO relative to hTG SP-A lines. Significant differences in viral antigens were observed among hTG SP-A mouse lines. (C): Lungs were collected from infected and sham mice on day 6 pi and viral load was determined by RT-qPCR quantification of the N gene. The horizontal dotted line shows the limit of detection (LOD) observed in sham controls. The data indicate higher viral copy number in KO mice compared to SP-A hTG mice. Significant differences in genome copies were observed among hTG SP-A mouse lines. (D): Plaque assay quantification of infectious virus in lungs collected on days 2, 4, and 6 pi. We observed peak viral load on day 2 pi. Significant differences in infectious virus titer were observed among hTG SP-A mouse lines on day 2, but not on day 4 and 6. \**P*< 0.05 \*\**P*< 0.01 (relative to sham), #<0.05, ##<0.01 (between two infected groups) from 3-4 independent experiments. Scale bars= 50µm. Symbols in (B -D) represent mean ± SEM of individual animals.

### Human SP-A Variants Differentially Modulate Innate and Adaptive Immune Gene Expressions in the Lungs of SARS-CoV-2 Infected Humanized Mice

Given that human SP-A can interact with both viral and host proteins to modulate immune responses, we assessed the role of SP-A in modulating immune gene expression post-viral challenge in this mouse model ^50^. We used a microarray-based system that allowed us to study the transcriptional changes of a focused panel of 84 innate and adaptive immune genes in the lungs from infected and sham groups. In the hierarchical clustering of genes illustrating gene-expression changes in infected vs mock-challenged mice, we observed that KO and 1A^0^ mice (these animals showed the most severe disease) had the greatest numbers of differentially expressed genes (DEGs) such as *Il18*, *NLRP3*, Jak2, *Stat3, Irak1* and *Il1r1* (Fig. 5A); reflecting a more enhanced activation of inflammatory genes in these groups. Of the 84 genes studied, 17 and 15 genes were significantly upregulated in infected KO and 1A^0^ relative to mock-challenged mice respectively (DEGs were ranked by >2.0-fold change, p<0.05) (Fig 5A). Extended Table 2 shows all the DEGs in infected and sham groups.

**Fig. 5.**
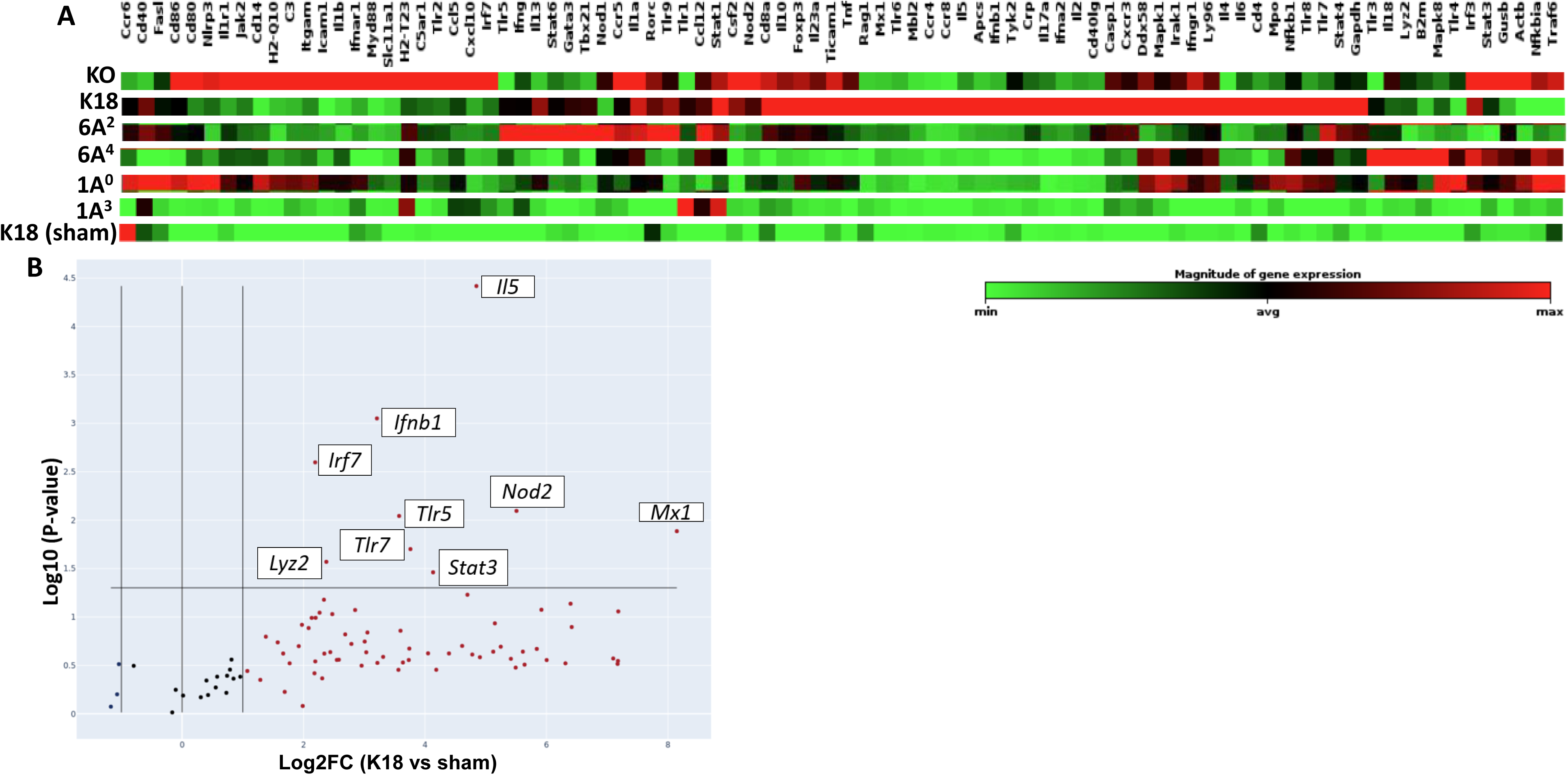
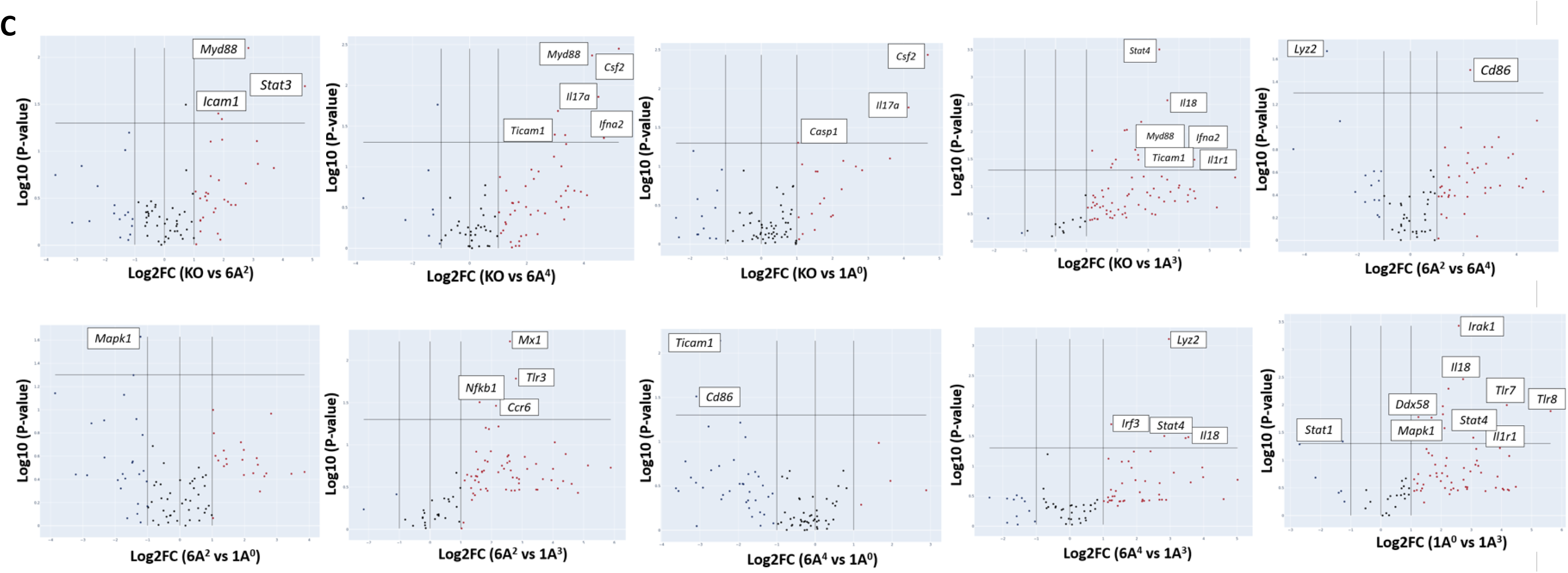
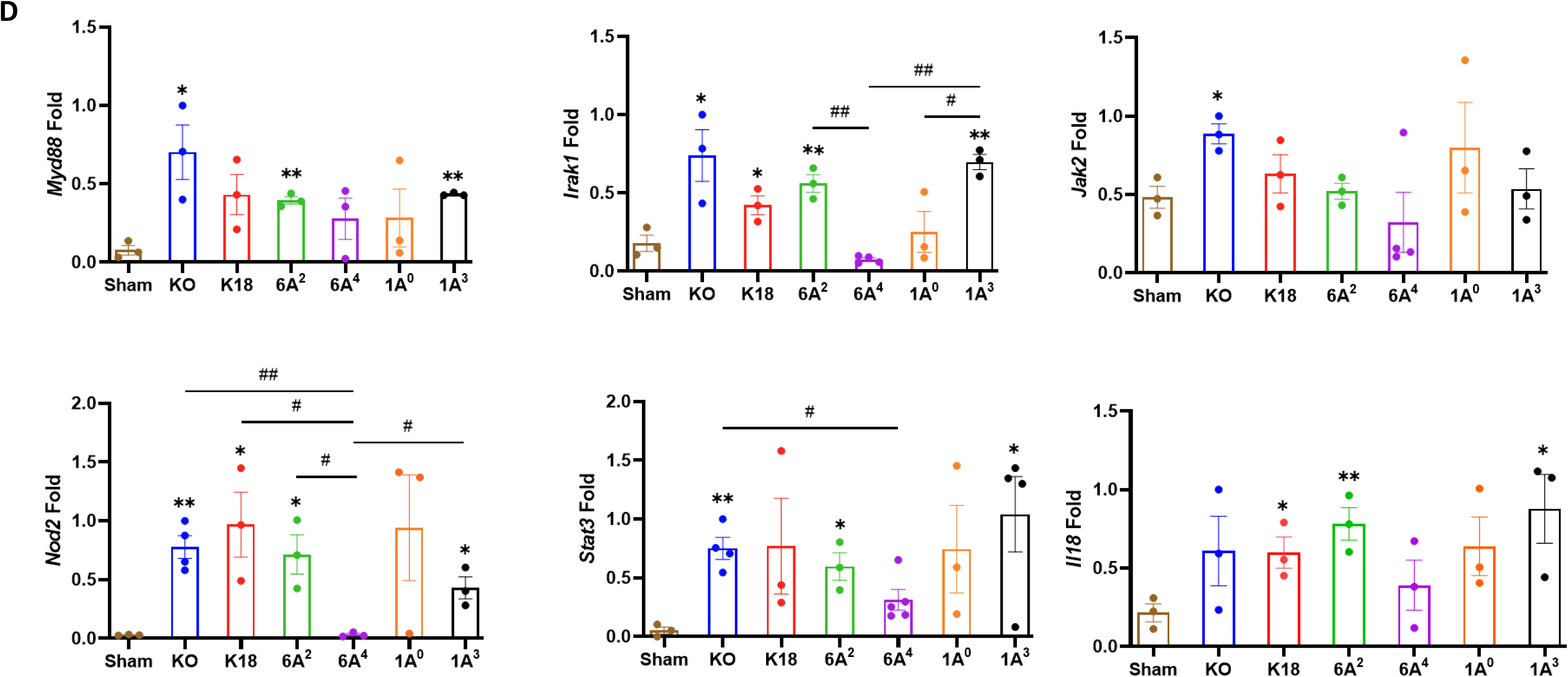
Differential transcriptional signatures in innate and adaptive immune genes in the lung of SARS-CoV-2 infected humanized mouse lines. A total of 84 innate and adaptive immune genes were examined in the lung tissues of six infected mouse lines, including sham mice. (A): Hierarchical clustering of genes illustrating changes in gene expression compared between infected mice and sham mice (n=3 mice/group). Of the 84 genes studied, 17 and 15 genes were significantly upregulated in infected KO and 1A^0^ mice relative to mock-challenged mice, respectively. (B): Volcano plots of differentially expressed genes (DEGs>2.0-fold change, p<0.05) between infected K18 mice relative to mock-challenged mice. (C): Volcano plots of some significant DEGs between two infected groups (Top panels: KO vs 6A^2^; KO vs 6A^4^; KO vs 1A^0^; KO vs 1A^3^; 6A^2^ vs 6A^4^. Bottom Panels: 6A^2^ vs 1A^0^; 6A^2^ vs 1A^3^; 6A^4^ vs 1A^0^; 6A^4^ vs 1A^3^ & 1A^0^ vs 1A^3^). The data revealed an upregulation in immune genes such as *MyD88*, *Stat3*, & *Icam1* in KO vs 6A^2^; *IL17a*, *Csf2* and *Casp1* in KO vs 1A^0^ and *Mapk1* in 6A^2^ vs 1A^0^ mice. (D): RT-qPCR validation of DEGs in SARS-CoV-2 infected hTG Mice. Results showed significant differences in several gene expressions between infected mice and sham, as well as among hTG SP-A mouse lines. \**P*< 0.05 \*\**P*< 0.01 (relative to sham), #<0.05, ##<0.01 (between two infected mouse groups).

The analysis further revealed a significant upregulation of Myeloid Differentiation Primary Response 88 (*Myd88*), a unique signaling adaptor protein that has been shown to activate inflammatory cytokines through the NF-kB and MAPK1 pathways following SARS-CoV-2 infection in KO mice compared to other lines (Fig 5B-C). Other significant genes upregulated in KO mice relative to SP-A hTG and K18 mice are *Mx1*, *Icam1*, *Stat3*, *Nod2*, *Casp1*, and *Mapk1*. The more DEGs involved in cytokine induction and macrophage activation such as *Mapk1, Irak1* and *Il18* observed in KO and 1A^0^ mice on day 6 pi reflects a more sustained activation of inflammatory genes and, thus, worse pathology in these lines. Further analysis revealed that the antiviral molecule, *Mx1* was significantly upregulated in K18 mice and appears to be a key innate host defense mediator in this line relative to all other groups as previously observed in critical COVID-19 patients ^51^.

The Volcano plots of the two SP-A1 variants revealed the upregulation of *Cd86*, a costimulatory molecule for T-cell mediated immunity activation, and a downregulation of *Lyz2* (a marker for macrophage activation) in infected 6A^2^ mice (moderate disease) vs 6A^4^ mice (severe disease), suggestive of the important role of T cell-mediated immune responses in virus clearance. In contrast with the low number of DEGs observed in SP-A1, SP-A2 genotype 1A^0^ (critical disease) had several significantly upregulated proinflammatory immune genes such as *Il18*, *Mapk1*, *Irak1*, *Nfkb1*, and *Ticam1* compared to 1A^3^ (moderate disease). Interestingly, *Mapk1* was the only significant DEG downregulated in infected 6A^2^ mice relative to 1A^0^ mice. In 1A^3^ mice with moderate disease, 2 of 4 upregulated genes relative to sham were involved in antiviral state (*Mx1* and *STAT1*), neutrophil/monocyte recruitment (*CCL2*), and anti-inflammatory response (*IL10*). Thus, mice with critically severe disease (KO and 1A^0^) had more significant upregulation of immune gene expression and the higher activation of these genes may have contributed to the severe morbidity and mortality observed in these groups. Thus, differences in the capacity of human SP-A genetic variants in inflammatory gene modulation in the context of SARS-CoV-2 infection could result in differential disease severity post-SARS-CoV-2 infection (Fig 5B-C). Furthermore, we validated by RT-qPCR a select group of essential DEGs in disease development and observed remarkable differences in their expression levels (Fig 5D).

### Human SP-A Variants Differentially Modulate Inflammatory Signaling Pathways in SARS-CoV-2 Infected Humanized Mice

We analyzed the DEGs using the Ingenuity Pathway Analysis (IPA) tool to identify the signaling pathways implicated in each hTG mouse line. The DEGs were defined as those genes with fold change ≥ 2 and p-value ≤ 0.01. Data analysis of the respective genes that mapped to a given canonical pathway obtained using a ratio of the number of significant DEGs to those that map to a given pathway revealed the upregulation of major inflammatory response pathways such as, “Pathogen-induced cytokine storm”, “neuroinflammation”, “role of pattern recognition receptors in recognition of bacteria and viruses”, “Trem1” and “macrophage classical activation” signaling in infected hTG lungs relative to sham lungs (Fig 6A). These findings support the robust induction of cytokines and disease development in infected mice compared to sham mice. A comparison of the pathways specifically upregulated in mice with the most severe mortality (KO) compared to those with moderate mortality (6A^2^) showed that most genes mapped to Trem1 (p-value = 1.26E-7) and IL33 (p-value = 1.76E-6) signaling pathways. IL-33 activation has been reported in advanced COVID-19, contributing to severe lung inflammation by activating several proinflammatory cytokines such as IL-6 and TNF-α through the MyD88-MAPK-NF-kB signaling axis ^52–54^. As shown in Fig 6A and B, other pathways identified in KO mice relative to SP-A hTG lines with moderate disease are those related to the activation of inflammatory responses as described above. Several unique pathways activated in 1A^0^ vs 1A^3^ are neuroinflammation (p-value = 9.87E-21), “Toll-like receptor” (p-value = 1.22E-19), “pathogen-induced cytokine storm” (p-value = 1.32E-17) signaling pathways, indicating that the severe mortality and ALI observed in KO and 1A^0^ mice (groups with the most severe disease) may be due to an over-activation of immune genes resulting in a dysregulated cytokine storm in these mouse groups compared to those carrying the 6A^2^ and 1A^3^ (moderate disease) variants.

**Fig. 6.**
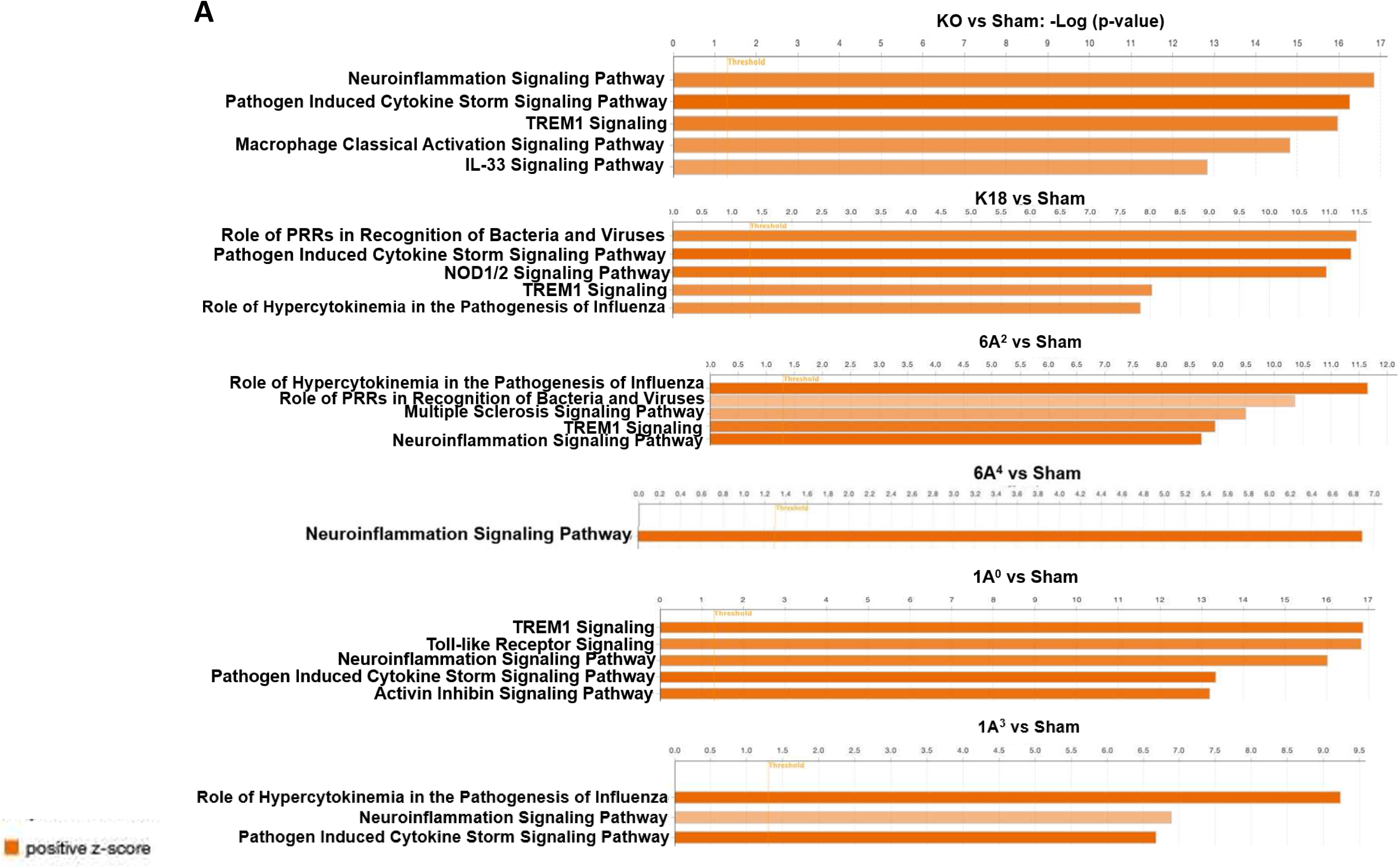
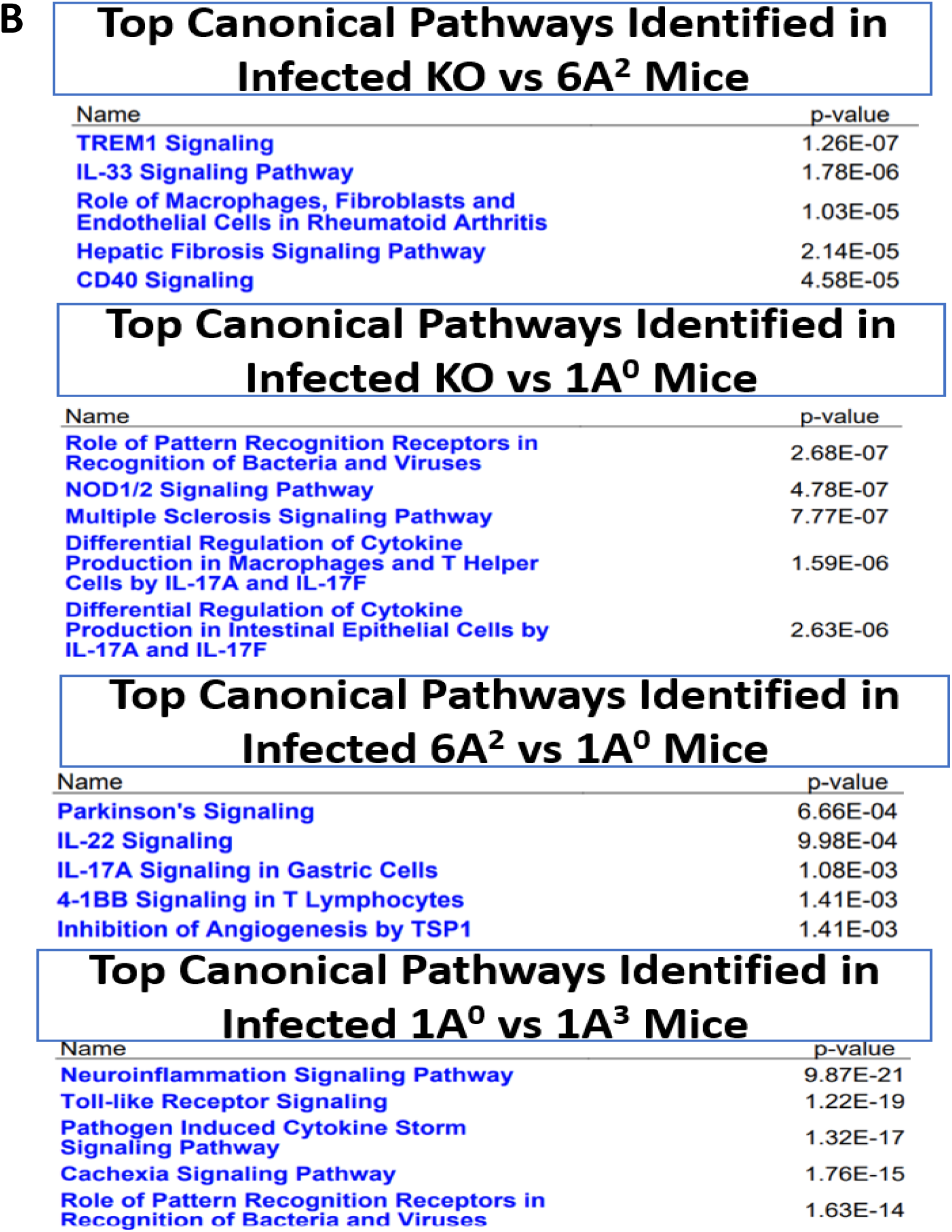
Top canonical pathways identified from functional analysis of DEGs in the lung of infected mouse lines. Top canonical pathways were identified using ingenuity pathway analysis (IPA). Canonical pathways were identified using only the significant up-and down-regulated genes from our microarray studies and ranked according to their –log (p-value) (Orange bars). Color intensity highlights the magnitude of fold change expressions. (A): Top one to five canonical pathways identified in SARS-CoV-2 infected double-hTG mice relative to sham mice. (B): Top five canonical pathways identified between two infected mouse groups: KO vs 6A^2^, KO vs 1A^0^, 6A^2^ vs 1A^0^, and 1A^0^ vs 1A^3^.

### Differential Cytokine Signatures in the Lungs of SARS-CoV-2 Infected Humanized Mice

Since the most severe COVID-19 groups (KO and 1A^0^) showed significant upregulation of multiple immune-related genes in the lungs, we evaluated the inflammatory signatures in mice carrying individual SP-A single-gene variants by quantifying the level of some cytokines in the lungs of infected and mock-challenged mice. We observed that IL-1β, IL-6, and IL-10 were significantly higher in KO, 1A^0^, and 6A^4^ compared to 6A^2^ and 1A^3^ mice (Fig 7A and B); but there was no significant difference in the levels of these cytokines in infected 6A^2^ and 1A^3^ mice relative to sham. The latter supports the role of elevated cytokines as major contributors to severe disease observed in the most severe groups. TNF-α was increased in all infected lines compared to sham, however, its level trended lower in 6A^2^ relative to other double-hTG lines. A Comparison of cytokine profiles between the two SP-A1 genotypes showed that IL-10 and IFN-γ were significantly higher in 6A^4^ mice (severe disease) compared to 6A^2^ mice (moderate disease). Data analysis of SP-A2 gene variants revealed higher levels of IL-1β, IL-6, and IL-10 in 1A^0^ mice (severe disease) vs 1A^3^ mice (moderate disease). These observations also correlated with the significant upregulation of immune genes in the lungs of KO mice and 1A^0^-carrying mice and consistent with cytokine profiling in COVID-19 patients which showed higher IL-6, IL-10, and TNF-α as correlates of severe COVID-19 ^1,55,56^.

**Fig. 7.**
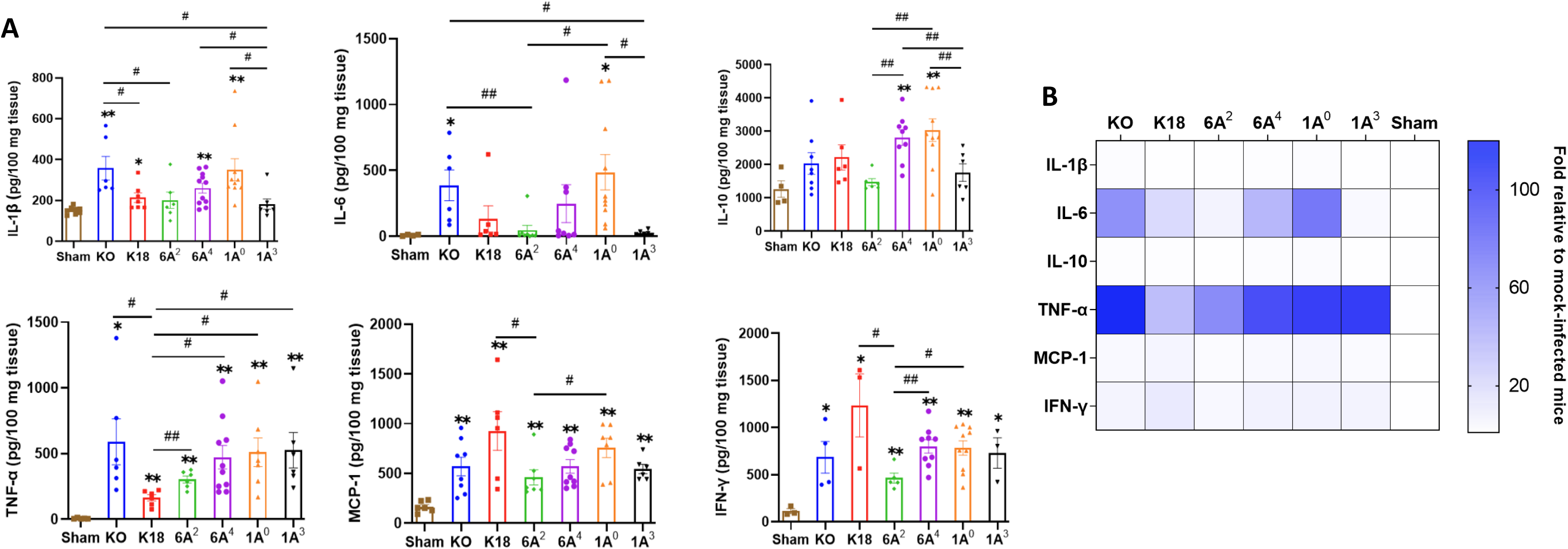
Differential cytokine expressions in the lungs of SARS-CoV-2 infected humanized mouse lines. (A): Bar graphs of absolute concentrations of cytokines and chemokines using ELISA in the lungs of infected and sham mice on day 6 pi. B: Heatmap representation of cytokine and chemokines levels in the lungs of SARS-CoV-2 infected mice normalized to levels in sham mice. Symbols in (A) represents mean ± SEM of individual animals. \**P*< 0.05 \*\**P*< 0.01 (relative to sham), #<0.05, ##<0.01 (between infected mouse groups), n= 6-10 mice/group.

### Differential Cytokine Signatures in Sera of SARS-CoV-2 Infected SP-A Humanized Mice

In severe COVID-19 patients, pulmonary and extrapulmonary manifestations such as acute kidney, intestinal, and brain injuries are observed ^55^. These extrapulmonary manifestations are often associated with increased levels of systemic cytokines and fatal SARS-CoV-2 infection ^55,57^. Therefore, we further measured the levels of 23 cytokines in the sera of SARS-CoV-2-infected and sham mice using a multiplex assay. Data analysis revealed a similar concentration of cytokines such as IL-1α, IL-2, GM-CSF, and IL-13 between infected and sham mice irrespective of the individual SP-A genotype (Fig 8A and B). Interestingly, as observed in human COVID-19 patients with fatal disease (KO and 1A^0^) and those with severe disease (6A^4^) had overall, significantly increased systemic levels of G-CSF, IL-6, IL-10, TNF-α, IL-12 (p40), IL-17A, CCL5, and CCL11 compared to those with moderate disease (6A^2^ and 1A^3^) ^55,58,59^. However, we found that serum levels of the T-cell associated cytokine (IFN-γ) were markedly elevated in mice carrying 6A^2^ and 1A^3^ variants (moderate disease) relative to KO and 1A^0^ (severe disease) mice. This finding is in line with previous reports on the importance of IFN-γ signaling and T-cell mediated immune responses in alleviating severe COVID-19 ^60,61^.

**Fig. 8.**
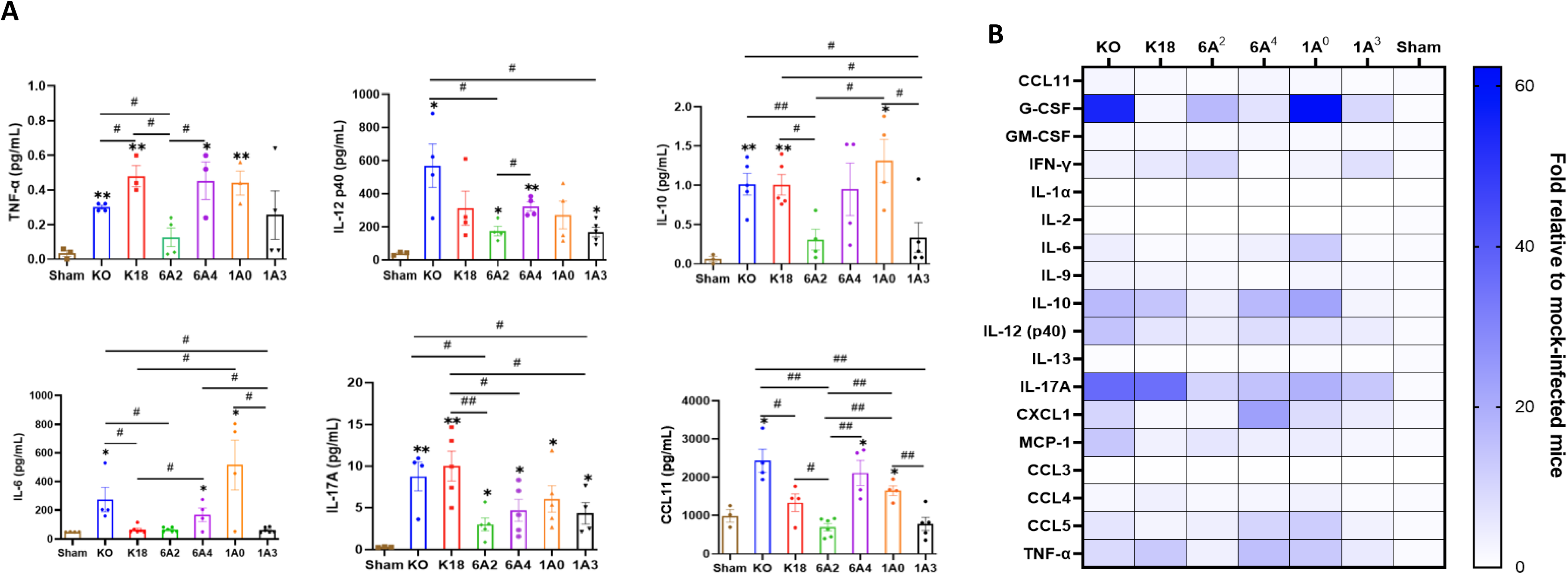
Differential cytokine signatures in sera of SARS-CoV-2 infected humanized mouse lines. A: Bar graphs of absolute levels of some cytokines obtained from multiplex assay in the sera of infected mice relative to sham on day 6 pi. B: Heatmap representation of cytokine and chemokines levels quantified using a multiplex platform in sera of SARS-CoV-2 infected mice on day 6 pi normalized to levels in sham mice. Symbols in (A) represents mean ± SEM of individual animals. \**P*< 0.05 \*\**P*< 0.01 (relative to sham), #<0.05, ##<0.01 (between infected mouse groups), n= 3-5 mice/group.

**Fig. 9.**
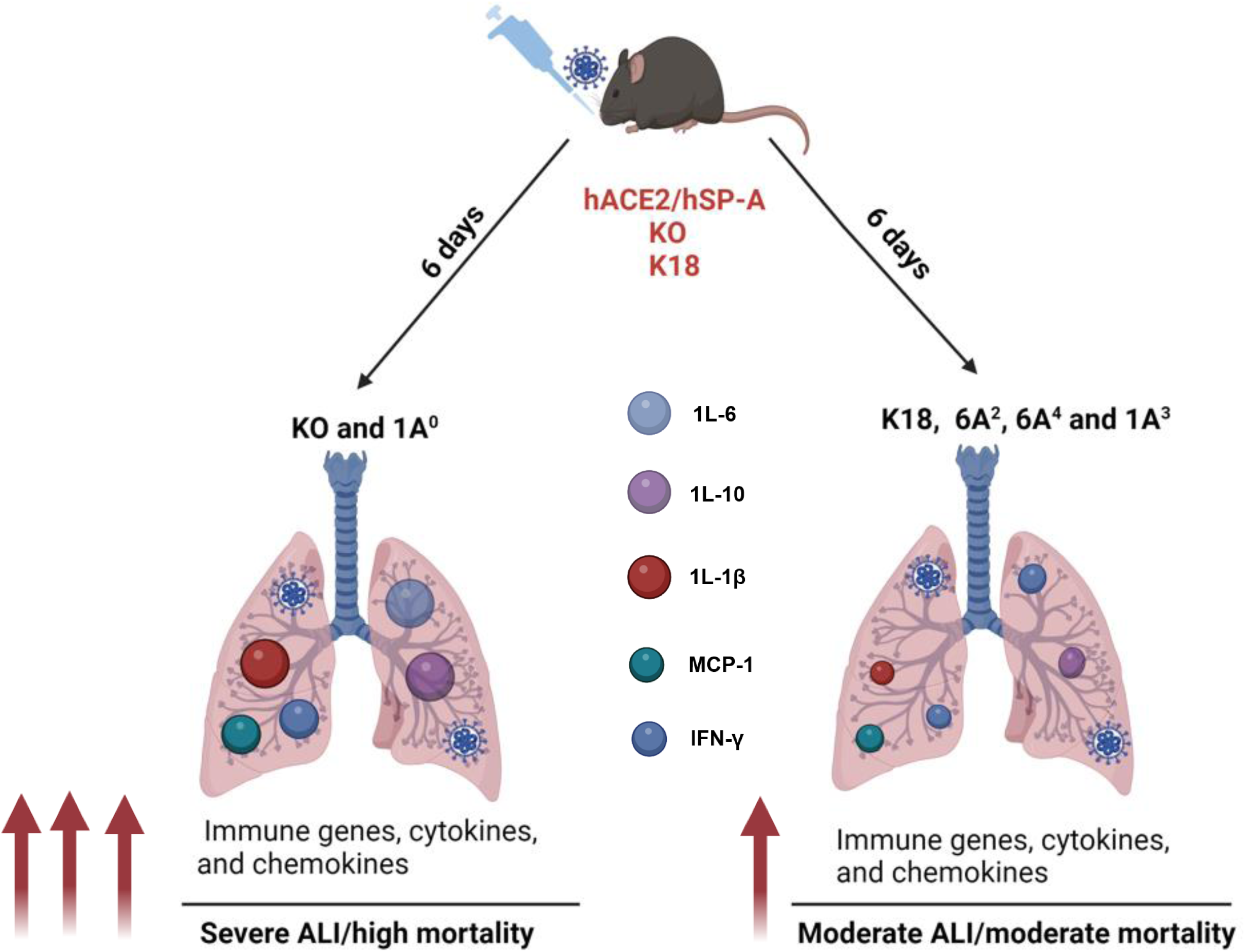
A model of the differential role of human SP-A genetic variants against SARS-CoV-2-induced ALI and mortality. SP-A variants differentially inhibit viral load in the lung after SARS-CoV-2 inoculation. ALI and disease severity are characterized by enhanced expressions of immune genes as well as higher levels of cytokines and chemokines in the lungs. Infected SP-A KO and 1A^0^ mice exhibited higher levels of innate and adaptive immune gene expression and elevated levels of pro-inflammatory cytokines and chemokines compared to infected K18, 6A^2^, 6A^4^ and 1A^3^ mice.

## Discussion

Besides identifying those susceptible to a given disease, the goal of precision medicine is to tailor treatment and prevention strategies using the unique gene set of the individual ^62^. Although most SARS-CoV-2 cases are mild, often requiring no hospitalization, some patients develop fatal respiratory distress in the absence of known risk factors like smoking and pre-existing medical conditions ^63–65^. In-depth GWAS analyses have linked these COVID-19 inter-individual variabilities to genetic variations in host proteins such as those of the blood group system, SARS-CoV-2 entry receptors, and innate and adaptive immune responses as drivers of severe disease observed among apparently healthy individuals ^34,35,66–70^.

As important as GWAS studies are at finding associations, pre-clinical studies are needed to determine the causal roles of these genes in human disease. Herein, we demonstrated that mice carrying single-gene variants of human SP-A1 and SP-A2 differentially modulate COVID-19 severity in response to SARS-CoV-2 infection. The results showed that (1) individual variants of human SP-A genes alleviate SARS-CoV-2 morbidity and mortality via their differential antiviral effects, (2) immune genes are differentially activated in mice carrying unique human SP-A variants, and (3) cytokine levels vary systemically and in the lungs in the mouse lines following SARS-CoV-2 infection and their levels directly correlate with severe disease. These observations mechanistically elucidated the roles of human SP-A genetic variants in the observed differences in COVID-19 symptomatology in the general population for the first time.

To decipher the innate immune roles of human SP-A genetic variants, we determined lung injury and animal mortality following SARS-CoV-2 infection and evaluated the established correlates of severe disease-viral load and inflammatory cytokine induction in mice ^49,57^. In this report, we observed two distinct patterns of COVID-19 severity in our hTG mouse lines, defined by survival rate and ALI severity, respectively. First, we observed that SP-A is important in preventing severe COVID-19 since SP-A deficient mice had the highest mortality rate and ALI scores compared to the other infected lines with either mouse SP-A (K18) or the respective human SP-A genes. This supports previous studies that observed higher viral titers in animals and patients with severe morbidity and mortality and reinforces the significant role of host innate pattern recognition molecules such as collectins in alleviating severe COVID-19 by inhibiting viral infectivity as was previously shown in the context of bacterial-induced pneumonia, RSV, flu, and recently, SARS-CoV-2 ^10,13,48,57,71–75^. Importantly, since the 6A^4^ and 1A^3^ mice had a more attenuated ALI, we speculate that there could be a small subset of individuals particularly protected from severe lung damage after SARS-CoV-2 infection ^76,77^. In addition, individuals predominantly carrying the 6A^2^ and 1A^3^ alleles may have better survival rates following infection than those carrying 1A^0^ and 6A^4^ alleles as previously observed with better lung function and mechanics in 1A^3^ *vs* 1A^0^ mice post-bacterial infection ^47^. The profound ALI and death in the SP-A2 (1A^0^) line was, however, intriguing and deviates from previous reports of high viral titers as drivers of severe disease given the significantly reduced viral titers observed in this line ^13,75^. The finding however aligns with a previous report of an association between the 1A^0^ variant and an increased risk of acute respiratory failure after IAV infection in humans and an animal model of pneumonia ^28,47^. Therefore, besides its antiviral function, we speculated that SP-A may modulate disease severity in response to SARS-CoV-2 infection via its other effector functions.

It has been established that COVID-19 is a cytokine-release syndrome caused by a dysregulated and over-activated immune response to infection. We speculate that while human surfactant protein (SP) genes play vital host defense functions against SARS-CoV-2 infectivity, their capacity to modulate inflammation in response to an infection may be even more crucial for disease modulation as was previously observed ^23,29,43,75,78^. Collectins such as SP-A, SP-D, and MBL largely serve as first line of defense against invading microbes by neutralizing, opsonizing, and facilitating microbial phagocytosis to enhance antigen presentation and stimulate the adaptive immune response ^79^. Importantly, from both immunological and biophysical standpoints, SP-A genetic variants differ in several instances: (1) the capacity for macrophages to phagocytose bacteria was higher in the presence of the 1A^0^ variant compared to 6A^2^; (2) the 1A^0^ variant was shown to activate inflammatory cytokine production more than 6A^2^ variant; (3) Structurally, SP-A2 gene-specific variants (1A^0^ and 1A^3^) are more stable and may better withstand oxidation by reactive oxygen radicals during inflammation compared to SP-A1 gene-specific variants (6A^2^ and 6A^4^); and (4) The proteomics and miRNAs of alveolar macrophages are different between SP-A1 and SP-A2 genes ^22,23,47,78^. Thus, the severe ALI and mortality observed in 1A^0^ mice may be due to some of the latter functional capacities of the SP-A2 (1A^0^) gene.

The transcriptomics data supported our hypothesis and revealed enhanced activation of immune genes in SP-A deficient and 1A^0^ mice compared to the other mouse lines. It appears that in the absence of SP-A, viral replication is unimpeded, resulting in an overstimulation and recruitment of immune cells, resulting in severe disease. However, the high mortality observed in the 1A^0^ line may be due to more viral aggregation, internalization by alveolar macrophages, enhanced innate immune signaling, and greater presentation of antigenic peptides in the lung environment to induce robust immune cell infiltration into the lungs. Insufficient, or delayed type 1 responses resulting in prolonged hyperinflammation, cellular influx, cell death, and lung fibrosis are some hallmarks of severe COVID-19. A delicate balance must thus be struck by innate and adaptive immune responses against the virus and modulation of these inflammatory responses to prevent sustained and hyperinflammatory tissue damage in the context of COVID-19. It is essential to state that our transcriptomic data were performed using samples collected 6 dpi, which is indicative of a sustained innate pro-inflammatory immune response in KO and 1A^0^ mice and consistent with persistent inflammation observed in severe patients ^80,81^.

T-cell-mediated immunity’s importance in preventing COVID-19 death was particularly highlighted with the upregulation of *Cd86,* a T-cell co-stimulatory molecule in 6A^2^ mice with moderate mortality compared to 6A^4^ mice with severe mortality. SARS-CoV-2, upon TLR-2 ligation, activates the Myd88 adaptor protein, subsequently signaling either through Mapk1 or NF-kB pathways ^82^. Interestingly, SP-A can activate NF-kB and induce cytokine production by directly interacting with TLR-2 and TLR-4 or by differentially regulating miRNA, which targets both proinflammatory and anti-inflammatory cytokine production ^83–86^. Moreover, SP-A was shown to reduce Mapk1 phosphorylation and activation ^87^. The downregulation of *Mapk1* in 6A^2^ mice relative to 1A^0^ is an interesting observation as it may impact cytokine production, which may have contributed to the higher survival rate recorded in mice carrying the 6A^2^ gene. Therefore, sequence changes in 6A^2^ could have caused functional differences in any of the latter mechanisms, impacting *Mapk1* activation and, potentially, cytokine levels in 6A^2^ mice.

A recurring canonical signaling pathway identified from our dataset is Trem1, an activating receptor mainly expressed in macrophages and neutrophils ^88^. Activation of Trem1 by soluble molecules, microbial ligands, inflammasome components, and cytokines through TLR ligation causes downstream activation of inflammatory cytokines and amplifies innate immune responses ^88,89^. Studies have shown that NLRP3 inhibition decreased Trem1 activation, resulting in diminished proinflammatory cytokine levels and disease severity ^89,90^. We further observed enrichment of Trem1 in infected KO compared to 6A^2^ mice, suggestive of the potential role of the 6A^2^ variant in attenuating severe disease by downregulating downstream inflammatory cytokine pathways. Further studies should address the underlying mechanism involved in the differential modulation of inflammatory responses via Trem1 signaling by SP-A variants. Do SP-A variants bind to Trem1 to interfere with subsequent activation by proinflammatory cytokines and viral structural proteins in myeloid cells ^88,90^? It remains to be elucidated whether SP-A variants attenuate inflammatory cytokine production by directly binding to Trem1 or by indirectly interacting with TLRs to reduce cytokine production and mitigate the positive feedback loop of cytokines on Trem1 activation.

To explain the differences in disease severity, we speculated that 1A^0^ mice may be better at inducing inflammatory responses than other SP-A genetic variants, given the high number of DEGs observed from our transcriptomic data. Our immunoassay of inflammatory cytokines in the lungs strongly supports this speculation where we observed higher levels of these cytokines in KO and 1A^0^ mice with the highest mortality compared to 6A^4^, 6A^2^, and 1A^3^ with high to moderate mortality rates (Mortality Rate: KO>1A^0^>6A^4^>1A^3^>K18>6A^2^). Certain cytokines directly correlate with severe disease or death, such as IFN-γ, TNF-α, IL-1β, IL-6, and IL-10 ^55,56^. The higher levels of IL-6 and IL-10 in KO and 1A^0^ mice in our study strongly correlate with previous clinical studies ^56,57,81^. We also speculate that variants 6A^2^ and 1A^3^ may be better than 1A^0^ at attenuating the levels of proinflammatory cytokines in macrophages stimulated with microbial or immune molecules as previously observed in functional studies using native SP-A and SP-A single-gene variants *in vitro* ^23,78,91^.

Cytokine expression and release in the context of SARS-CoV-2 infection is not only associated with ALI/ARDS but also with the development of multiple organ dysfunction (MOD) in COVID-19 patients ^55^. MOD has been linked with broad ACE2 expression, active viral replication in extrapulmonary organs, and high levels of systemic inflammatory cytokines ^92^. We therefore compared the levels of inflammatory cytokines in sera of individual hTG mice and observed that 6A^2^ and 1A^3^ mice with significantly higher survival rates had remarkably reduced levels of most cytokines except IFN-γ, a major inducer of cell-mediated immune responses, which are vital for clearing virally infected cells. Indeed, the higher systemic levels of IFN-γ observed in 6A^2^ mice support the double-edge sword theory of IFN-γ in COVID-19 pathology in which robust levels in blood may limit virus spread to other tissues, but increased levels in lungs may drive lung pathology as observed in the most severe mouse groups ^61,93^. This finding warrants further studies on the role of human SP-A genetic variants in modulating extrapulmonary disease manifestations of COVID-19.

SP-A exhibits functional and quantitative differences that are influenced by the unique SP-A haplotype, sex, and age of an individual, which ultimately influences health outcomes following infection due to transcriptomic and proteomic differences ^46,47,83^. One study showed enhanced miRNA activation in male mice, significantly impacting signaling molecules and proinflammatory cytokine induction ^83^. We therefore infected mice of similar ages in this study and observed that ALI severity and mortality trended higher in male K18, 6A^4^, and 1A^0^ compared to female mice. However, due to sample size limitation, a definitive contribution of sex in SP-A function cannot be ascertained and future studies should delineate the intricate interplay of age, sex, and the unique SP-A variant in the context of SARS-CoV-2 infection. Viral strain/variant differences in pathogenicity have been observed in animals and humans, so we speculate that the observed disease in the mouse groups may vary if SARS-CoV-2 variants other than Delta are used ^94^.

Changes in the levels of pulmonary surfactant proteins (SP-A and SP-D) and MBL have been shown to correlate with severe disease ^95^. Our previous data revealed a significant reduction in SP-A levels among severe COVID-19 patients while a dysregulation in SP genes was observed in SARS-CoV-2 infected lung tissues ^10,33^. The decrease in collectin levels post-primary infection should be further addressed in a co-infection model and emphasizes the need to develop lung surfactant treatments for people particularly predisposed to severe COVID-19.

For the first time, we demonstrated that human SP-A genetic variants differentially attenuate ALI and affect survival rates post SARS-CoV-2 infection, by reducing viral burden, and differentially modulating viral-induced inflammatory responses. Here, we have demonstrated that compared to SP-A2 (1A^0^), mice carrying the SP-A1 gene-specific variant (6A^2^) have the highest survival rate, followed by SP-A2 (1A^3^) and SP-A1 (6A^4^) and survival rate inversely correlated with lung cytokine levels. SP-A genetic variants (particularly 6A^2^ and 1A^3^) seem more capable of attenuating the robust induction of cytokines observed in severe COVID-19 and SP-A gene variants are protective (albeit differentially) in response to SARS-CoV-2 infection by distinctively activating immune genes and signaling pathways. We, therefore, speculate that morbidity and mortality observed in some infected individuals may be explained by the unique and predominant SP-A genotypes in such individuals. Understanding the contributions of human SP-A genetic variants in response to SARS-CoV-2 infection is vital to developing novel gene/variant-specific surfactant-based therapies for COVID-19 patients.

## Conflict of Interest Statement

The authors confirm that no competing interest exists.

Author Contributions: G.W., H.J., S.T., and I.B.J. contributed to the conception and study design. I.B.J., A.O. L., and E.R. performed ELISA and multiplex assays. I.B.J. E.R., S.T and A.O.L., S.S.M., E.M.K. and Q.M contributed to infectious virus titration and mouse challenge studies. I.B.J., P.T.M, H.J., H.F., and G.W. contributed to infection assays and data analysis. I.B.J. P.T.M., H.F., and G.W. drafted the first version of the manuscript. All authors read and approved the manuscript.

## ACKNOWLEDGMENTS

This work was supported by NIH R01HL136706, R21 AI171574, and the National Science Foundation research award (1722630) (to GW) and by the 1R01AI148446-01A1 (to HJ). We thank the staff of the Upstate Vector Biocontainment Laboratory (BSL-3) for their technical support and SUNY Upstate Pathology Research Core (SUNY SPORE) for their histological services. We would like to thank Dr. Joanna Floros for her generous support with the biomaterials used for this project and her scientific perspective on this study. Vero E6 cells were obtained through BEI Resources, NIAID, and NIH.

## Notes

### Competing Interest Statement

The authors have declared no competing interest.

