## Supplemental data for "Differential Immunoregulation by Human Surfactant Protein A Variants Determines Severity of SARS-CoV-2-induced Lung Disease"

**Running title:** Impact of Human SP-A Variants in SARS-CoV-2-Induced ALI

**Corresponding author:**

Guirong Wang, PhD (Dr. rer. Nat.)

Department of Surgery, UH Room 8715

SUNY Upstate Medical University

750 E Adams St,

Syracuse, NY, 13210, USA

**Conflict of Interest Statement**

The authors confirm that no competing interest exists.

Author Contributions: G.W., H.J., S.T., and I.B.J. contributed to the conception and study design. I.B.J., A.O. L., and E.R. performed ELISA and multiplex assays. I.B.J. E.R., S.T and A.O.L., S.S.M., E.M.K. and Q.M contributed to infectious virus titration and mouse challenge studies. I.B.J., P.T.M, H.J., H.F., and G.W. contributed to infection assays and data analysis. I.B.J. P.T.M., H.F., and G.W. drafted the first version of the manuscript. All authors read and approved the manuscript.

**Results**

**Extended Table 1: Primer List For RT-qPCR Validation**

| **Gene (Mouse)** | Forward primer (5'- 3') | Reverse primer (5'- 3') |
| --- | --- | --- |
| **Il18 201/202** | AAGTGCCAGTGAACCCCAGACCA | CACAGAGAGGGTCACAGCCAGTCC |
| **MyD88 203/204** | TCCGACCGTGACGTCCT | ACCATGCGGCGACACC |
| **Irak1** | TCCACCAAGCAGTCAAGCC | AAACCACCCTCTCCAATCC |
| **NOD2** | GCTGTCTTGGGATGTGCT | GGATGAAGGGAGTGAGTGTC |
| **Gapdh** | CCAATACGGCCAAATCC | CCAATACGGCCAAATCCG |
| **Stat3** | GGGCATTTTTATGGCTTTCAAT | GTTAACCCAGGCACACAGACTTC |
| **Jak2** | AGGCGACGGGAACAAGATGT | AGGCCATTCCCATCTAGAGC |

**Extended Fig.1. Differential lung injury after SARS-CoV-2 infection of humanized mouse lines**

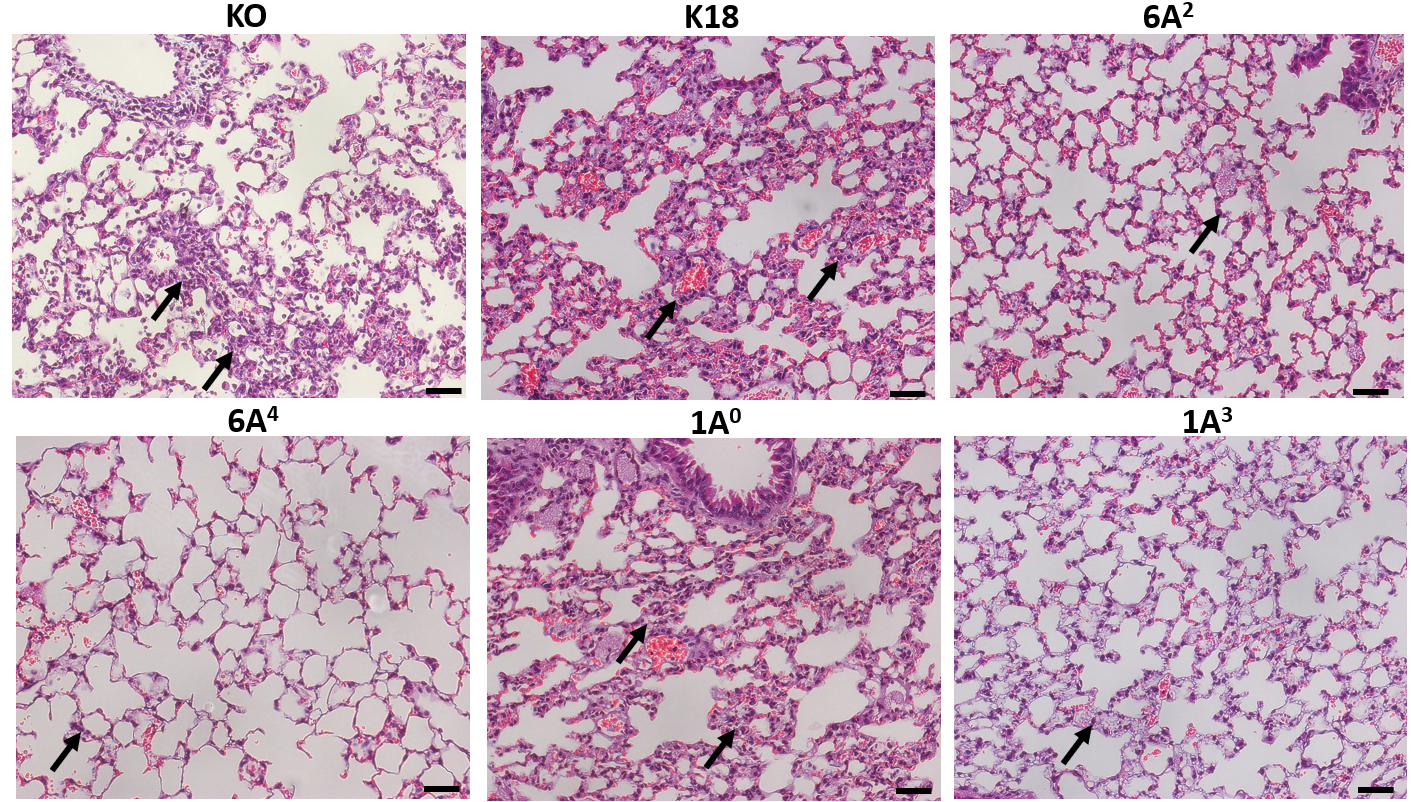

**Extended Table 2**

| Differentially Expressed Genes (Infected KO vs Sham) | | |
| --- | --- | --- |
| Gene Symbol | **Fold Regulation** | **P-value** |
| *Casp1* | 5.79 | 0.014756 |
| *Cd5* | 8.91 | 0.046393 |
| *Icam1* | 7.69 | 0.008094 |
| *Ifngr1* | 4.39 | 0.044389 |
| *Il17a* | 16.48 | 0.022416 |
| *Il18* | 19.19 | 0.004625 |
| *Il1r1* | 7.97 | 0.020679 |
| *Irak1* | 3.92 | 0.016144 |
| *Jak2* | 11.98 | 0.022278 |
| *Lyz2* | 5.63 | 0.019558 |
| *Myd88* | 6.40 | 0.010672 |
| *Nod2* | 17.93 | 0.023428 |
| *Stat1* | 6.28 | 0.017607 |
| *Stat3* | 12.27 | 0.014362 |
| *Actb* | 4.13 | 0.041631 |
| *B2m* | 6.17 | 0.030071 |
| *Gusb* | 3.54 | 0.023738 |
| Differentially Expressed Genes (Infected K18 vs Sham) | | |
| *Ifnb1* | 28.76 | 0.000038 |
| *Il5* | 45.44 | 0.008027 |
| *Irf7* | 17.54 | 0.034547 |
| *Lyz2* | 4.57 | 0.002529 |
| *Mx1* | 283.34 | 0.012998 |
| *Nod2* | 11.91 | 0.009057 |
| *Stat3* | 5.19 | 0.026936 |
| *Tlr5* | 9.24 | 0.000889 |
| *Tlr7* | 13.55 | 0.019901 |
| Differentially Expressed Genes (Infected 6A^2^ vs Sham) | | |
| *Casp1* | 5.76 | 0.035678 |
| *Ifng* | 118.07 | 0.016573 |
| *II10* | 76.39 | 0.019744 |
| *Mx1* | 74.29 | 0.002725 |
| *Nfkb1* | 2.71 | 0.049131 |
| *Tlr3* | 3.99 | 0.026076 |
| Differentially Expressed Genes (Infected 6A^4^ vs Sham) | | |
| *Il18* | 30.81 | 0.029020 |
| *Lyz2* | 12.26 | 0.000422 |
| *Stat1* | 4.30 | 0.013537 |
| *Tlr5* | 6.31 | 0.024732 |
| *B2m* | 16.77 | 0.038125 |
| Differentially Expressed Genes (Infected 1A^0^ vs Sham) | | |
| *Ccl5* | 4.78 | 0.018746 |
| *Cd86* | 9.82 | 0.040562 |
| *Ddx58* | 4.14 | 0.007225 |
| *Il18* | 18.55 | 0.002340 |
| *Il1r1* | 5.43 | 0.036498 |
| *Irak1* | 4.72 | 0.001273 |
| *Ly96* | 5.71 | 0.034474 |
| *Mapk1* | 4.18 | 0.027030 |
| *Mapk8* | 4.44 | 0.011781 |
| *Nfkb1* | 3.82 | 0.033683 |
| *Stat3* | 6.64 | 0.042926 |
| *Tlr7* | 9.23 | 0.023362 |
| *Tlr8* | 41.35 | 0.012885 |
| *Gapdh* | 10.85 | 0.002370 |
| *Gusb* | 2.61 | 0.036120 |
| Differentially Expressed Genes (Infected 1A^3^ vs Sham) | | |
| *Ccl2* | 16.05 | 0.025912 |
| *Il10* | 21.73 | 0.049830 |
| *Il18* | 2.80 | 0.036129 |
| *Mx1* | 12.25 | 0.022571 |
| *Stat1* | 6.81 | 0.006183 |
| Differentially Expressed Genes (KO vs K18) | | |
| *Icam1* | 7.61 | 0.018423 |
| *Myd88* | 13.16 | 0.004582 |
| *Il5* | -4.54 | 0.026129 |
| *Mx1* | -9.55 | 0.020986 |
| Differentially Expressed Genes (KO vs 6A^2^) | | |
| *Icam1* | 3.53 | 0.039577 |
| *Myd88* | 7.14 | 0.007947 |
| *Stat3* | 26.81 | 0.020250 |
| *Actb* | 3.84 | 0.045359 |
| Differentially Expressed Genes (KO vs 6A^4^) | | |
| *Csf2* | 37.43 | 0.003549 |
| *Ifna2* | 26.07 | 0.044254 |
| *Il17a* | 22.79 | 0.013839 |
| *Myd88* | 19.56 | 0.004296 |
| *Nod2* | 7.90 | 0.040216 |
| *Ticam1* | 8.54 | 0.020631 |
| *Tyk2* | 10.17 | 0.040750 |
| *Lyz2* | -2.18 | 0.017263 |
| Differentially Expressed Genes (KO vs 1A^0^) | | |
| *Casp1* | 2.05 | 0.049517 |
| *Csf2* | 25.50 | 0.003665 |
| *Il17a* | 17.56 | 0.017414 |
| Differentially Expressed Genes (KO vs 1A^3^) | | |
| *Gata3* | 12.35 | 0.002657 |
| *Icam1* | 6.43 | 0.018416 |
| *Ifna2* | 22.71 | 0.032396 |
| *Ifngr1* | 4.73 | 0.009398 |
| *Il17a* | 7.60 | 0.021143 |
| *Il18* | 6.85 | 0.006584 |
| *Il1r1* | 17.28 | 0.014299 |
| *Irak1* | 4.97 | 0.009201 |
| *Jak2* | 6.44 | 0.032430 |
| *Lyz2* | 3.58 | 0.038574 |
| *Mapk1* | 2.28 | 0.022174 |
| *Myd88* | 14.54 | 0.016160 |
| *Nod2* | 9.60 | 0.028490 |
| *Stat3* | 6.00 | 0.028496 |
| *Stat4* | 10.29 | 0.000313 |
| *Ticam1* | 6.39 | 0.026506 |
| *Actb* | 3.48 | 0.044980 |
| *Gusb* | 3.87 | 0.032193 |
| Differentially Expressed Genes (K18 vs 6A^2^) | | |
| *Mx1* | 3.81 | 0.034390 |
| Differentially Expressed Genes (K18 vs 6A^4^) | | |
| *Csf2* | 29.29 | 0.006292 |
| *Ifnb1* | 61.15 | 0.000059 |
| *Il5* | 20.37 | 0.008441 |
| *Mx1* | 9.81 | 0.038784 |
| *Nod2* | 5.25 | 0.039805 |
| *Lyz2* | -2.68 | 0.002301 |
| Differentially Expressed Genes (K18 vs 1A^0^) | | |
| *Csf2* | 19.95 | 0.006556 |
| *Ifnb1* | 8.06 | 0.003828 |
| *Il5* | 7.32 | 0.016924 |
| *Mx1* | 8.69 | 0.025430 |
| Differentially Expressed Genes (K18 vs 1A^3^) | | |
| *Ifnb1* | 20.78 | 0.000018 |
| *Il5* | 49.38 | 0.007167 |
| *Irf3* | 2.22 | 0.039937 |
| *Lyz2* | 2.90 | 0.015772 |
| *Mx1* | 23.13 | 0.014983 |
| *Nod2* | 6.38 | 0.013982 |
| *Tlr5* | 3.27 | 0.002488 |
| *Tlr7* | 26.55 | 0.012512 |
| Differentially Expressed Genes (6A^2^ vs 6A^4^) | | |
| *Cd86* | 4.78 | 0.031320 |
| *Lyz2* | -8.85 | 0.021367 |
| Differentially Expressed Genes (6A^2^ vs 1A^0^) | | |
| *Mapk1* | -2.32 | 0.023491 |
| Differentially Expressed Genes (6A^2^ vs 1A^3^) | | |
| *Ccr6* | 4.43 | 0.034447 |
| *Mx1* | 6.06 | 0.005957 |
| *Nfkb1* | 3.06 | 0.031248 |
| *Tlr3* | 6.94 | 0.016428 |
| Differentially Expressed Genes (6A^4^ vs 1A^0^) | | |
| *Cd86* | -8.54 | 0.030853 |
| *Ticam1* | -5.56 | 0.007244 |
| Differentially Expressed Genes (6A^4^ vs 1A^3^) | | |
| *Gata3* | 11.67 | 0.033026 |
| *Il18* | 11.00 | 0.034283 |
| *Irf3* | 2.36 | 0.020149 |
| *Lyz2* | 7.80 | 0.000772 |
| *Stat4* | 7.09 | 0.031649 |
| Differentially Expressed Genes (1A^0^ vs 1A^3^) | | |
| *Cd86* | 8.37 | 0.039116 |
| *Ddx58* | 2.37 | 0.016583 |
| *Fasl* | 5.44 | 0.012656 |
| *Il18* | 6.62 | 0.003422 |
| *Il1r1* | 11.77 | 0.020857 |
| *Irak1* | 5.99 | 0.000372 |
| *Ly96* | 6.50 | 0.025717 |
| *Mapk1* | 3.20 | 0.016987 |
| *Mapk8* | 4.15 | 0.010562 |
| *Nfkb1* | 4.32 | 0.026328 |
| *Stat4* | 9.48 | 0.025254 |
| *Ticam1* | 4.16 | 0.014790 |
| *Tlr7* | 18.08 | 0.010077 |
| *Tlr8* | 49.47 | 0.012963 |
| *Gapdh* | 4.73 | 0.005095 |
| *Stat1* | -2.40 | 0.045731 |
